# Mettl1-dependent m^7^G tRNA modification is essential for maintaining spermatogenesis and fertility in *Drosophila melanogaster*

**DOI:** 10.1101/2023.09.04.555845

**Authors:** Shunya Kaneko, Keita Miyoshi, Kotaro Tomuro, Makoto Terauchi, Shu Kondo, Naoki Tani, Kei-Ichiro Ishiguro, Atsushi Toyoda, Hideki Noguchi, Shintaro Iwasaki, Kuniaki Saito

## Abstract

N^7^-methylguanosine (m^7^G) in the variable loop region of tRNA is catalyzed by METTL1/WDR4 heterodimer and stabilizes target tRNA. Here, we reveal essential functions of Mettl1 in *Drosophila* fertility. Knockout of Mettl1 (Mettl1-KO) lost the elongated spermatids and mature sperm, which was fully rescued by a Mettl1-transgene expression, but not a catalytic-dead Mettl1 transgene. This demonstrates that Mettl1-dependent m^7^G is required for spermatogenesis. Mettl1-KO resulted in a loss of m^7^G modification on a subset of tRNAs and a decreased level of tRNA expression. Strikingly, overexpression of the translational elongation factor, EF1α1, which can compete with the rapid tRNA decay (RTD) pathway in *S. cerevisiae*, significantly counteracted the sterility of Mettl1-KO males, supporting a critical role of m^7^G modification of tRNAs in spermatogenesis. Ribosome profiling showed that Mettl1-KO led to the ribosome stalling at codons decoded by tRNAs that were reduced in expression. Mettl1-KO also significantly reduced the translation efficiency of genes involved in elongated spermatid formation and sperm stability. These findings reveal a developmental role for m^7^G tRNA modifications and indicate that m^7^G modification-dependent tRNA stability differs among tissues.

## Introduction

Posttranscriptional epigenetic modifications of RNA are together known as the “epitranscriptome” and are being revealed to have diverse and widespread functions in gene regulation. To date, over 170 RNA modifications have been identified, approximately 80% of which are found in tRNAs^1,2^. Chemical modifications regulate tRNA structure, dynamics and interactions with other molecules and result in altered translation^3,4^. While some tRNA modifications play a fundamental role in cell proliferation, an increasing number of studies show that aberrant tRNA modification can result in severe developmental defects and human diseases, indicating that tRNA modifications play roles in specific tissues during animal development^4–7^.

N7-methylguanosine (m^7^G) is an RNA modification observed in mRNA caps^8^ and at internal positions within tRNAs, rRNAs, mRNAs and miRNAs^9–15^. m^7^G in tRNAs is widely conserved in prokaryotes, eukaryotes and some archaea^16^. In eukaryotes, the RNA methyltransferase-like 1 (METTL1) catalyzes the addition of m^7^G at internal sites of tRNAs, mRNAs, and miRNAs^10,12,15,17^. METTL1 interacts with the non-catalytic subunit WD repeat domain 4 (WDR4)^17–21^, and both are widely conserved from yeast to mammals. m^7^G modification occurs at the 46th guanosine in the variable loop region of tRNAs, within the conserved motif, “RAG<u>G</u>U” (R = A or G, <u>G</u> represents m^7^G)^10^. In *Saccharomyces cerevisiae*, transfer RNA methyltransferase 8 (Trm8), a homolog of METTL1, introduces the m^7^G modification onto tRNAs, such as tRNA ValAAC, and promotes tRNA stability and cell growth at higher temperatures^17,18,22^. Hypomodified tRNAs are degraded by the 5′–3′ exonucleases, Rat1 and Xrn1, which form the rapid tRNA decay (RTD) pathway^22–24^. Rat1 and Xrn1 are highly conserved among animals, indicating that a similar degradation system is present in animals^25^. Indeed, METTL1-depletion in mammalian cells leads to decreased steady-state levels of several tRNAs that are targets of METTL1, retardation of cell growth in mammalian cells^26,27^ and impaired translation of genes associated with neural differentiation^10^. Conversely, increased levels of METTL1 result in increased m^7^G modification of a specific tRNA, (such as tRNA ArgTCT), tRNA stabilization, cell cycle acceleration through enhanced translation of cell cycle related genes, such as *CDK6* and *CDK8*, and promotion of oncogenicity^27^. The primary *let-7e* miRNA is also an RNA methylated by METTL1^12^. In cultured human cells, m^7^G affects the secondary structure of primary *let-7e* and promotes the processing into precursor *let-7e*^12^. In mRNAs, internal m^7^G modifications are recognized by Quaking proteins, which enable their transport into stress granules. This modulates mRNA stability and the cellular stress response in mammals^28^. These studies have revealed the role of m^7^G at the single cell level; however, its roles at tissue and organ levels during animal development are less well characterized.

Recent clinical studies have revealed that variations in *METTL1* and *WDR4* causes microcephalic primordial dwarfism, Galloway-Mowat syndrome^29–31^, multiple sclerosis^32,33^, and male fertility^34^, indicating a potential link between internal m^7^G regulation and pathological characteristics^35^. Furthermore, knockout of *WDR4*/*mWh* in mice results in embryonic lethality^36^. These observations indicate that the m^7^G modification machinery is critical for mammalian development; however, it is unclear whether m^7^G modification alone is responsible for the above developmental defects. Indeed, WDR4 physically interacts with proteins other than METTL1. WDR4 is proposed to be involved in genome stability through interaction with Flap Endonuclease 1^36^ and in cerebellar development and locomotion through interaction with Atrgap17^37^. Therefore, it is not known whether the developmental disorders caused by METTL1 or WDR4 variants are directly linked to defective m^7^G modification.

In *Drosophila melanogaster*, *Wuho (Wh)*, the homolog of WDR4, is required for gonadal development^38,39^. *Wh* mutants show male sterility with arrested spermatogenesis in males and semi-sterile phenotypes with abnormal egg chamber and decreased germline stem cell number in females^38,39^. Wh is considered to control ovary function through interaction with partner proteins, including Mei-P26, Nanos, and Bgcn^38^. However, it remains to be addressed whether Wh associates with the *Drosophila* METTL1 ortholog and if the fertility defects in *Wh* mutants are caused by a loss of m^7^G modification or via defective association with interactors of Wh other than Mettl1.

Here, we demonstrated that candidate gene 4045 (CG4045)/ Mettl1 is required for *Drosophila* male and female fertility. Mettl1 associates with Wh *in vivo* and *in vitro* and catalyzes m^7^G-modification on a subset of tRNAs in a Wh-dependent mechanism. Depletion of Mettl1 results in a loss of m^7^G modification and decreased stability of nearly all, but not all, of the tRNAs targeted by Mettl1. Importantly, overexpression of the translational elongation factor, eEF1α1, which presumably competes with the RTD pathway, rescued Mettl1-KO male infertility, indicating that m^7^G-mediated tRNA stability is essential for male fertility. Consistent with the decrease in tRNA abundance, we found increased occurrence of ribosome pausing and subsequently collisions of trailing ribosomes in Mettl1-KO testis at the codons decoded by the reduced tRNAs. Our work further reveals the developmental role of Mettl1-dependent m^7^G in multicellular organisms. We propose that the developmental consequence of Mettl1-KO arises via a combination of the effects of impaired m^7^G-mediated tRNA stabilization, the RTD pathway, and competitors of the RTD pathway, such as eEF1α1.

## Results

### Mettl1, the Drosophila ortholog of the m^7^G-modifier, METTL1, is essential for *Drosophila* fertility

We studied the physiological functions of Mettl1 in *Drosophila melanogaster*. The *Drosophila* gene, CG4045 (hereafter referred to as Mettl1), is an ortholog of mammalian METTL1 and contains an RNA methyltransferase-like domain that is highly conserved across animals (Fig. 1a, Supplementary Fig. 1a)^40^. To generate a Mettl1-knock out (KO) strain using the CRISPR/Cas9 system, a guide RNA (gRNA) was selected to target the N-terminal part of the methyltransferase-like domain (Fig. 1a). Two different transgenic strains harboring either a 1 nucleotide deletion (*Mettl1^KO1^*) or a 23-nucleotide deletion with an 8-nucleotides insertions (*Mettl1^KO2^*) were generated, both of which lose a large part of the methyltransferase-like domain because of frame-shift (Fig. 1a, Supplementary Fig. 1b). To check the expression of Mettl1 and the Mettl1-interacting partner, Wh, mouse-monoclonal antibodies were generated. Western blot analyses showed a strong ∼32kDa Mettl1 signal in the heterozygous ovaries and the control (*yw*) testes (Fig. 1b, c). In contrast, this signal was undetectable in the *Mettl1^KO1^* and *Mettl1^KO2^*female ovaries and *Mettl1^KO1^* testes (Fig. 1b, c, Supplementary Fig 1c), indicating that Mettl1 was absent in Mettl1-KO animals. Notably, a Wh signal was still observed in Mettl1-KO flies (Fig. 1b), indicating that Wh stability is unaffected by the absence of its heterodimer partner, Mettl1.

**Figure 1.**
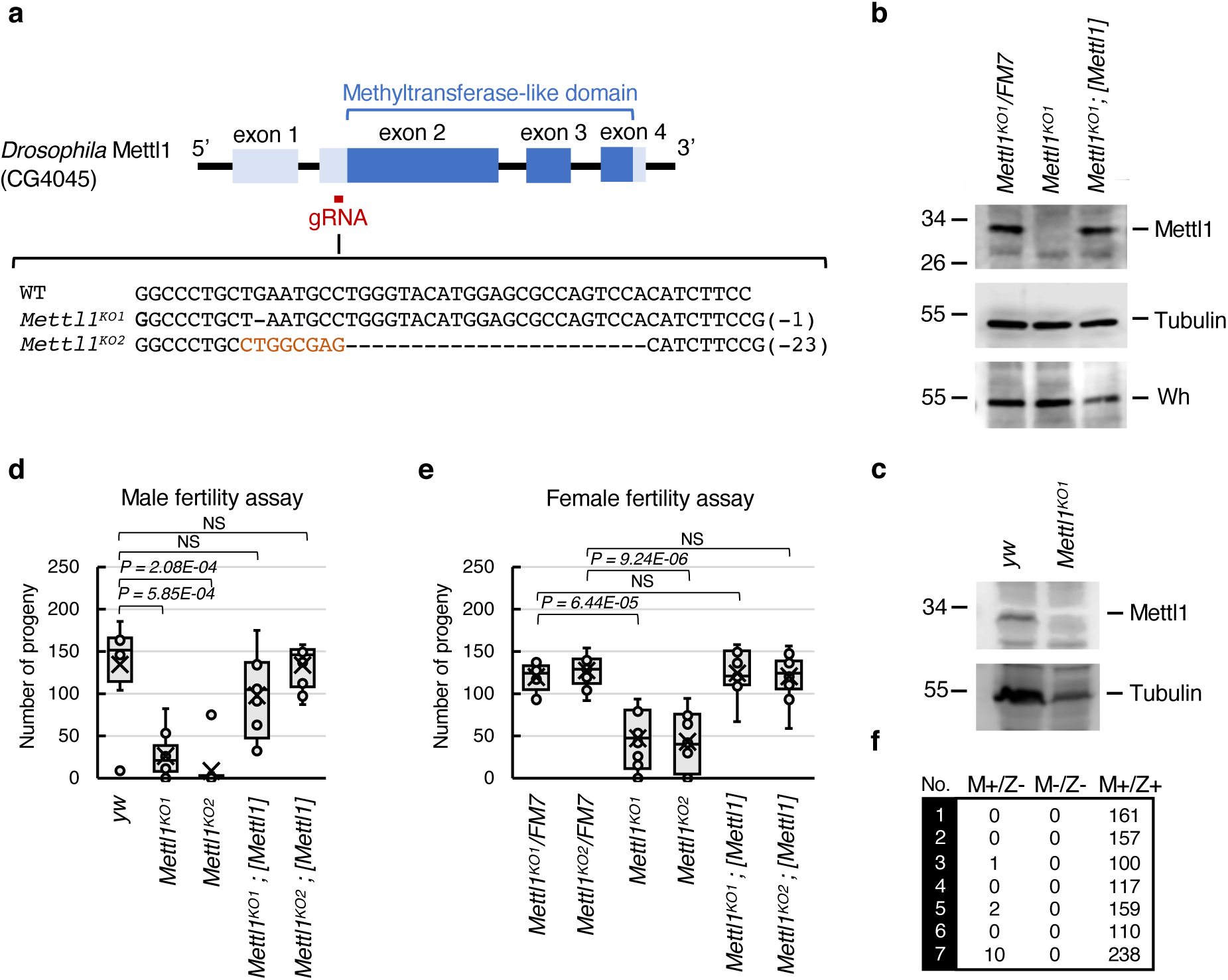
Mettl1 is important for maintaining fertility. **(a)** Schematic diagram of the genomic structure of Mettl1 and a gRNA targeting Mettl1 exon 2 for CRISPR-mediated generation of Mettl1-KO lines. Mettl1 sequences of the WT (*yw*) and the Mettl1-KO mutants (*Mettl1^KO1^* and *Mettl1^KO2^*). **(b)** Western blot showing Mettl1 and Wh expression in *Drosophila* ovaries. **(c)** Western blot showing Mettl1 expression in *Drosophila* testes. **(d)** Male fertility assay at 25℃ using WT (*yw*), Mettl1-KO (*Mettl1^KO1^*, *Mettl1^KO2^*) and Mettl1-rescue (*Mettl1^KO1^*; *[Mettl1]*, *Mettl1^KO2^*; *[Mettl1]*) males. **(e)** Female fertility assay at 25℃ using control groups (*Mettl1^KO1^/ FM7, Mettl1^KO2^/ FM7*), Mettl1-KO groups (*Mettl1^KO1^, Mettl1^KO2^*) and Mettl1-Rescue groups (*Mettl1^KO1^*; *[Mettl1]*, *Mettl1^KO2^*; *[Mettl1]*) females. FM7 is a balancer chromosome. **(f)** Male fertility assay at 25℃ using (*yw* ; m+/z+), and zygotic Mettl1-KO (*Mettl1^KO1^* ; m+/z–), maternal and zygotic Mettl1-KO (*Mettl1^KO1^* ; m–/z–) males. In fertility assay, box plots represent maximum, median and minimum values with outliers. Upper and lower edges of boxes represent third quartile and first quartile, respectively **(d, e)**. Significant differences calculated by two-tailed Student’s t-test were indicated on graph. NS means not significant.

### *Mettl1^KO1^* and *Mettl1^KO2^* flies appeared to develop normally to the adult stage

However, we observed that mutant males and females had significantly reduced fertility (Fig. 1d, e, Supplementary Fig. 1d). Mutant males produced one tenth and mutant females one third of the progeny produced by the control (Fig. 1d, e, Supplementary Fig. 1d). Consistent with the reduced fertility of Mettl1-KO females, the size of the Mettl1-KO ovaries was significantly reduced (Supplementary Fig. 1e). To confirm whether this is caused by loss of Mettl1 function, we introduced the Mettl1 genomic region into the second chromosome. The *attp40[Mettl1]* transgene rescued the fertility and ovary size to the levels of control groups (Fig. 1d, e, Supplementary Fig. 1e). Moreover, to evaluate the maternal contribution to fertility, we generated maternal and zygotic mutant males. Small numbers of offspring (less than 10) were detected in three of seven crosses from zygotic mutant males, whereas no offspring were produced by maternal and zygotic mutant males, indicating a maternal contribution of Mettl1 (Fig. 1f). Taken together, these data uncovered a critical role of Mettl1 in the male fertility.

### Mettl1 depletion impairs the formation of elongated spermatids and results in the loss of mature sperm

*Drosophila* spermatogenesis involves germ cells communicating with somatic cells, proceeding through mitotic and meiotic cell division, differentiating into elongating spermatids, and forming bundle structures in the testis. Each step in this process is tightly regulated (Fig. 2a) ^41^. To study the expression patterns of Mettl1 in the testis, we generated a transgenic strain carrying C-terminally Flag-tagged Mettl1, *[Mettl1-Flag]* (Fig. 2b). We observed Flag-tag signals in almost all testicular cells, indicating that Mettl1 is ubiquitously expressed in the various gonadal cell types (Fig. 2b). Wh-GFP shows a similar ubiquitous expression pattern through spermatogenesis, indicating the presence of the Mettl1/Wh heterodimer in all testicular cells^39^. To study whether spermatogenesis was affected by Mettl1-KO, we assessed mature sperm generation. We found no mature sperm in the seminal vesicles of Mettl1-KO testes (Fig. 2c), which is consistent with the decreased fertility of Mettl1-KO males (Fig. 1d). To examine the spermatogenesis defects in more detail, we studied the expression of the germ cell marker proteins, Aubergine, a marker of germline cells, and Boule, a *Drosophila* orthologue of the vertebrate DAZl, a marker of 32- and 64-cell cysts and elongating spermatids within *Drosophila* testes^42^. Aubergine was observed in a wild-type pattern within Mettl1-KO testes (Supplementary Fig. 2a). Moreover, Boule was also detected in a wild-type pattern in 32- and 64-cell cysts within Mettl1-KO testes (Fig. 2d, Supplementary Fig. 2b). Boule-positive cysts defect to elongation were accumulated near seminal vesicle region (Fig. 2d). These indicate that the loss of Mettl1 did not affect meiosis but may affect elongations step of spermatogenesis. Furthermore, we observed no elongated spermatids (Fig. 2e). We then examined the distribution of the acetylated microtubules (MTs), a marker of elongated spermatids^43^. Most of Mettl1-KO testes (23/31) showed a decreased acetylated MT signal (indicating an elongation defect, Fig. 2f), The remainder had no elongated spermatids (8/31, Fig. 2f). Together, these results indicate that the elongated spermatid formation and the later stages of spermatogenesis are severely impaired in Mettl1-KO testes.

**Figure 2.**
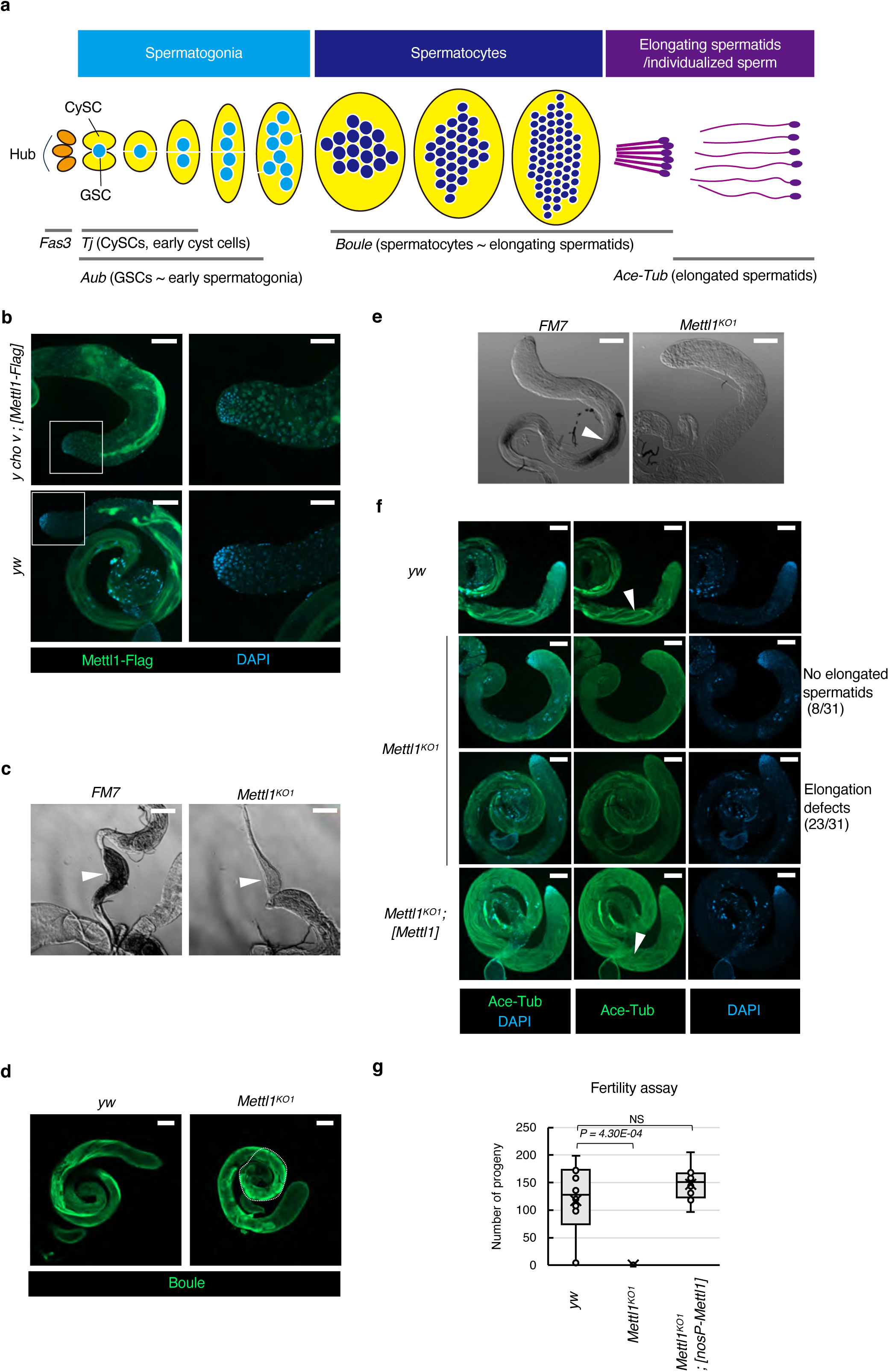
Loss of Mettl1 results in absence of mature sperm in testis. **(a)** Schematic diagram of *Drosophila* spermatogenesis. The genes indicated below are markers used to identify each stage of spermatogenesis. (GSC; Germline Stem Cell, CySC; Cyst Stem Cell) (**b)** Conformation of *[Mettl1-Flag]* expression in testis by immunostaining. *yw* is a control. (**c)** Seminal vesicle (arrowheads) from control (*FM7*) and Mettl1-KO (*Mettl1^KO1^*) males. (**d)** Representative images of Boule localization in testes from WT (*yw*), Mettl1-KO (*Mettl1^KO1^*) adult males. Testes were stained for Boule (green). (**e)** Differential interference microscopy image of the spermatid bundles (arrowheads) in *FM7* (control) and *Mettl1^KO1^* testes. (**f)** Acetylated-Tub localization shows elongated spermatids in WT (*yw*) and Mettl1-rescue (*Mettl1^KO1^; [Mettl1]*) testes but not in Mettl1-KO (*Mettl1^KO1^*) testes. (**g)** Number of progeny gained from male fertility assay using WT (*yw*), Mettl1-KO (*Mettl1 ^KO1^*), Mettl1-KO with Mettl1 transgene driven by Nanos promoter (*Mettl1^KO1^; [nosP-Mettl1]*). Box plots represent maximum, median and minimum values with outliers. Upper and lower edges of boxes represent third quartile and first quartile, respectively. Significant differences calculated by two-tailed Student’s t-test were indicated on graph. NS means not significant. Scale bars of **b**(left), **c**, **d**, **e**, and **f** = 100 μm, and **b**(right) = 50 μm.

We observed no change in overall morphology or in the distribution of somatic niche cells (hub cells) or somatic cyst cells in Mettl1-KO testes, indicating the requirement for Mettl1 in testicular germ cells (Supplementary Fig. 2c). Mettl1 is still detected in the carcasses of males after removal of the gonads (Supplementary Fig. 2d), indicating that Mettl1 may play a critical role in elongated spermatid formation in tissues other than the testis. To directly examine whether Mettl1 acts in a cell-autonomous or non-cell autonomous manner, we prepared a Mettl1-transgene driven by a germ-specific Nanos promoter and conducted a fertility assay. The Nanos-Mettl1-transgene *attp40[nosP-Mettl1]* fully rescued the sterile phenotype of Mettl1-KO males (Fig. 2g). Moreover, somatic cell-specific knockdown (KD) of Mettl1 did not affect fertility (data not shown). Taking these findings together, we concluded that Mettl1 plays a critical role in elongating spermatid formation via a cell-autonomous mechanism.

### Catalytic activity of Mettl1 is required for *Drosophila* fertility

Although mammalian METTL1 and its yeast ortholog, Trm8, catalyze the m^7^G modification on tRNAs, it is not known whether *Drosophila* Mettl1 possesses this activity^10,17^. Given that METTL1 acts as the catalytic partner in a heterodimer with WDR4^10,17,20,21^, we investigated the complex formation in flies. We conducted immunoprecipitation from the *[Mettl1-Flag]* ovaries or Oregon R wild-type ovaries using two different buffer conditions and performed liquid chromatography tandem mass spectrometry (LC-MS/MS) analysis of the samples (Fig. 3a, Supplementary Fig. 3a, b). Mettl1-Flag co-immunoprecipitated with Wh irrespective of buffer conditions used in the immunoprecipitaion, indicating that the Mettl1/Wh heterodimer is formed *in vivo*, as in other organisms (Fig. 3a, Supplementary Fig. 3a, b). Notably, we did not detect Mei-P26, Nanos or Bgcn proteins, which has been reported to associate with Wh^38^, indicating that Wh makes a distinct protein complex either with Mettl1 or with those proteins. To confirm the direct interaction, we purified MBP-Mettl1 and GST-Wh from *Eschericia coli* (*E. coli*) and performed *in vitro* GST-pull down assays (Fig. 3b). MBP-Mettl1 associated with GST-Wh but not with the GST moiety (Fig. 3b).

**Figure 3.**
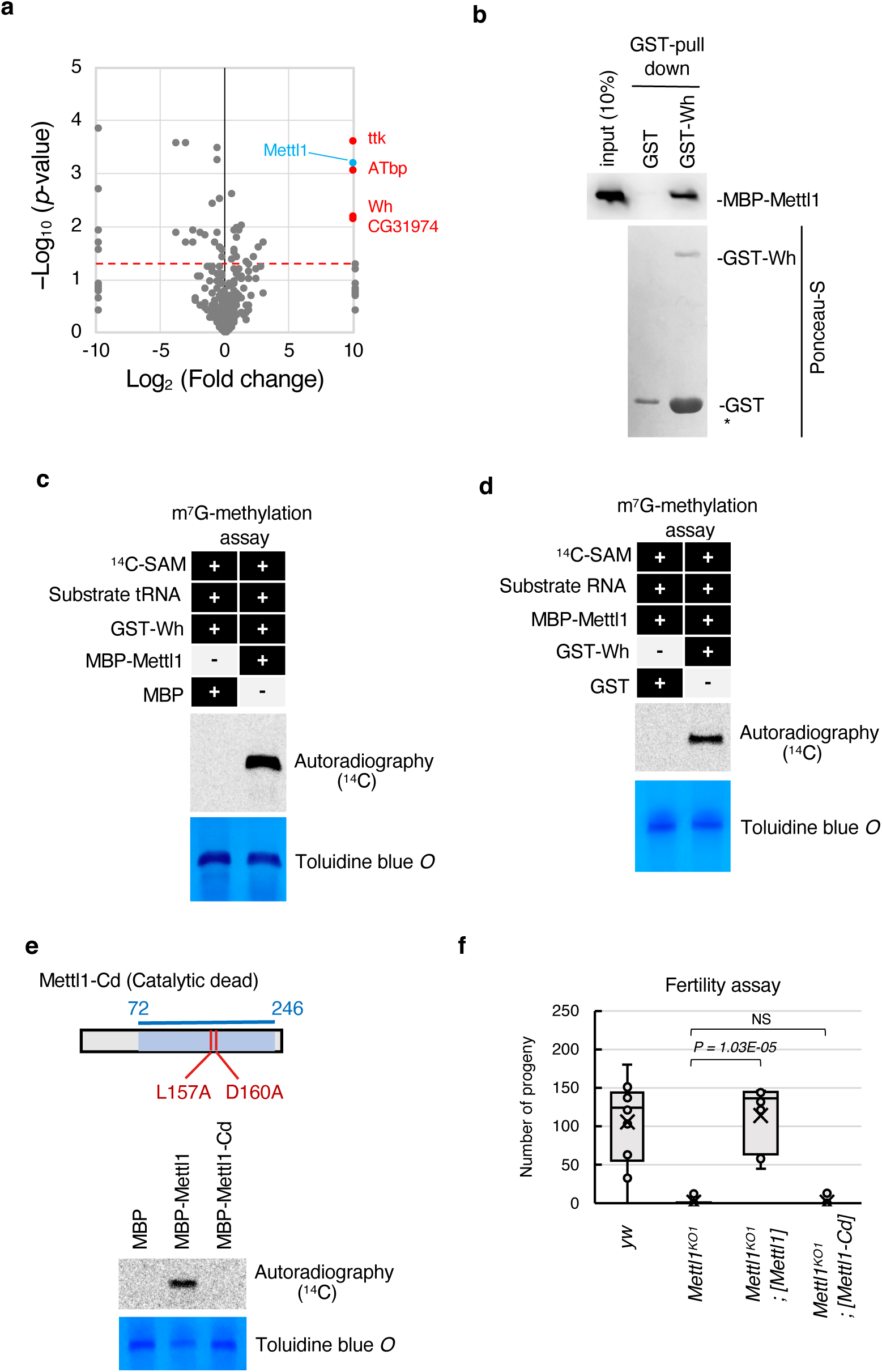
Catalytic activity of *Drosophila* Mettl1 is required for fertility. **(a)** Volcano plot showing enrichment rates and significance levels of each protein as log2 fold change (anti-Flag M2/negative control) versus negative log 10 of the fisher’s exact test p value. Four red dots represent Mettl1 interactors, and blue dot represents Mettl1 in HEPES-NP40 buffer, respectively. (**b)** GST-pulldown assays showing direct association between *Drosophila* Mettl1 and Wh. MBP-Mettl1 and GST-Wh were treated as prey and bait. GST was used as a negative control. (**c, d, e)** *In vitro* m^7^G modification methylation assay. tRNA TrpCCA is a tRNA substrate of m^7^G. The m^7^G containing tRNAs were labelled with 14C. Toluidine blue O staining was done to visualize RNAs. **c** Confirmation of *Drosophila* Mettl1 requirement in m^7^G methylation. This experiment used recombinant *Drosophila* Mettl1/Wh and MBP (negative control). (**d)** Confirmation of *Drosophila* Wh requirement in m^7^G methylation. This experiment used recombinant *Drosophila* Mettl1/Wh and GST (negative control). (**e)** Confirmation of requirement of amino acid residues that important of methylation activity of Mettl1 m^7^G methylation. This experiment used recombinant *Drosophila* Mettl1/Wh, Mettl1 harboring L157A and D160A mutation and MBP (negative control). (**f)** Male fertility assay at 25℃ using WT (*yw*), Mettl1-KO (*Mettl1^KO1^*) and Mettl1-rescue (*Mettl1^KO1^ ; [Mettl1]*) and catalytic dead Mettl1-rescue (*Mettl1^KO1^; [Mettl1-Cd]*) males. Box plots represent maximum, median and minimum values with outliers. Upper and lower edges of boxes represent third quartile and first quartile, respectively. Significant differences calculated by two-tailed Student’s t-test were indicated on graph. NS means not significant.

We then conducted *in vitro* m^7^G methylation assays using the above recombinant proteins with a synthesized tRNA and a shortened tRNA fragment designed according to a previous study^44^, both of which include the RAGGU motif, a conserved target sequence of eukaryotic METTL1 (Fig. 3c, Supplementary Fig. 3c). *Drosophila* Mettl1 required Wh for tRNA methylation, consistent with a recently proposed structural model of m^7^G methylation (Fig. 3d)^20,21^. Mettl1 did not introduce m^7^G to a target RNA in which the 4^th^ guanosine of the RAGGU motif was converted to cytosine, indicating the *Drosophila* Mettl1/Wh methylates target tRNAs in a similar manner to that of METTL1/WDR4 in other eukaryotes (Supplementary Fig. 3c)^10^. To create a catalytic dead mutant of Mettl1, L157A and D160A mutations (Mettl1-Cd: Cd), both of which are located in the methylation consensus motif, were engineered (Supplementary Fig. 1a). We confirmed that these mutations abolished methylation activity of Mettl1 *in vitro* (Fig. 3e).

This finding led us to test the requirement of Mettl1 catalytic activity for fertility. We prepared a catalytic dead Mettl1-transgene *[Mettl1-Cd]*, harboring L157A and D160A mutations in Mettl1 (Fig. 3f) and conducted rescue experiments of the fertility assay (Fig. 3f). The sterile phenotype of *Mettl1^KO1^*was rescued by the wild-type Mettl1-transgene *[Mettl1]*, but not by *[Mettl1-Cd]*, indicating that male fertility is dependent on the catalytic activity of Mettl1 (Fig. 3f).

### *Drosophila* Mettl1 mediates m^7^G modification of tRNA

To understand the molecular mechanism underlying the Mettl1-KO fertility defects, we surveyed m^7^G modification sites on tRNAs because tRNAs are efficiently methylated by METTL1 in mammalian cells^10^. We performed an m^7^G-site specific cleavage assay using sodium borohydride (NaBH_4_) and aniline^45,46^. The resulting 3′ fragments were detected by northern blotting using specific tRNA probes (Fig. 4a, b). We detected the 3′ fragments in the presence of Mettl1 but not in Mettl1-KO for tRNA isolated from testes (Fig. 4b) and ovaries (Supplementary Fig. 4a), supporting Mettl1 deposition of m^7^G onto tRNAs in both tissues (Fig. 4b, Supplementary Fig. 4a). Moreover, the 3′ fragments from the same tRNA were lost in Wh mutant testis (Fig. 4c), consistent with the collaborated role of Wh for m^7^G modification on tRNA (Fig. 4c, Fig. 3d).

**Figure 4.**
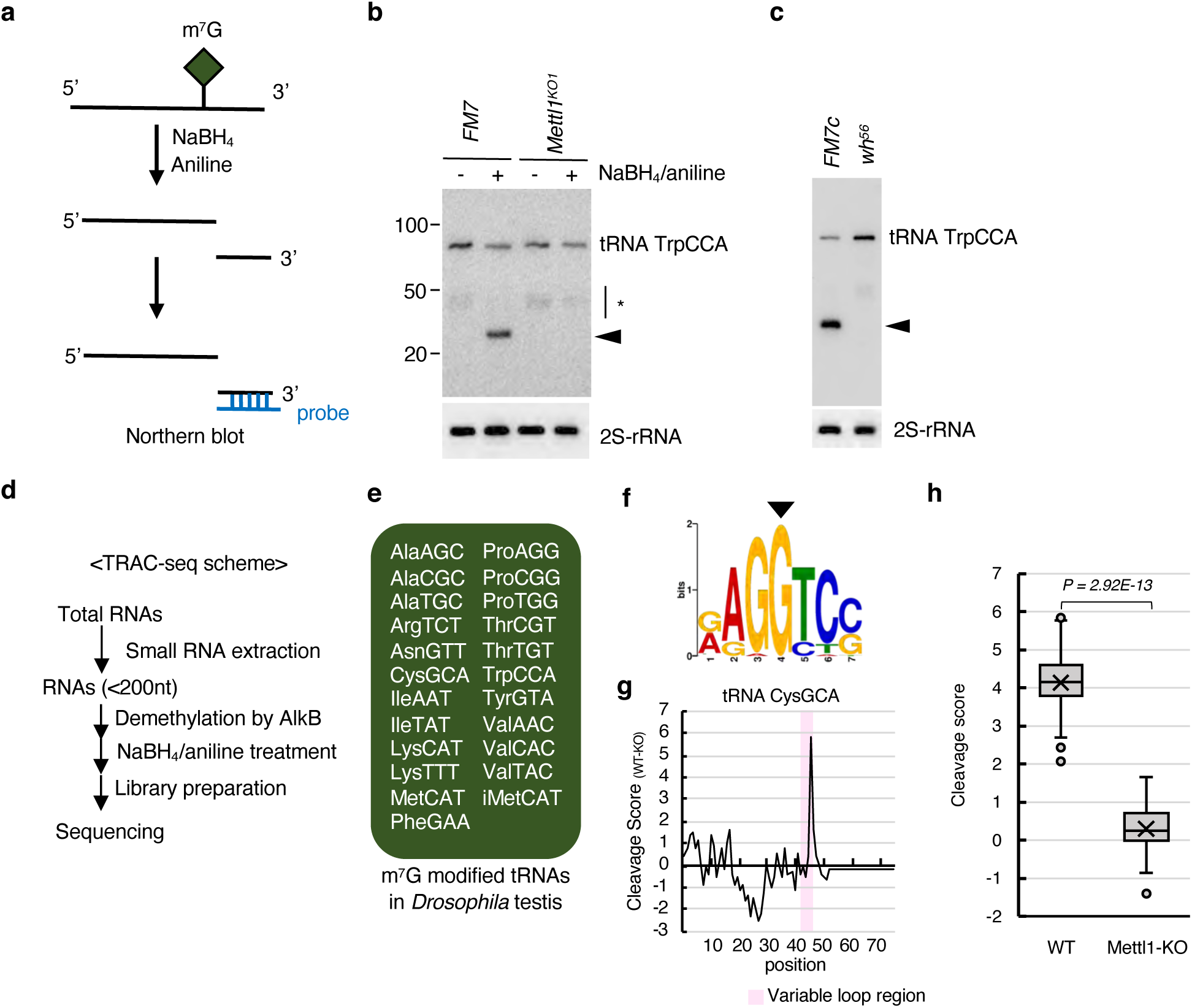
*Drosophila* Mettl1 methylates tRNA in gonad. **(a)** Schematic diagram of m^7^G-site specific reduction and cleavage. Cleaved 3′ fragments were detected by northern blotting. (**b)** Northern blot of the chemically-treated total RNAs from control (*FM7*) and Mettl1-KO (*Mettl1^KO1^*) testes. The probe was designed at the 3′ end of TrpCCA. **(c)** Northern blot of the chemical treated total RNAs from WT (*FM7c*) and Wh-KO (*Wh^56^*) testis. Probe was designed at around 3′ end of tRNA TrpCCA. **(d)** TRAC-seq scheme using *Drosophila* gonads. (**e)** Mettl1-dependent m^7^G modified tRNAs identified in testes by TRAC-seq. (**f)** Sequence motif of m^7^G modification sites identified in testes by the TRAC-seq. The arrowhead corresponds to the m^7^G site. (**g)** Representative plot of the cleavage score (difference between cleavage score of WT and Mettl1-KO) of tRNA CysGCA (Cys-GCA-2) between WT and Mettl1-KO, which showed the highest score difference by TRAC-seq. Pink shade represents variable loop region of tRNA CysGCA. (**h)** Quantitative comparison of cleavage scores (CleavageScore) of identified m^7^G-modified tRNAs between WT and Mettl1-KO. Significant differences calculated by Mann-Whitney’s U-test were indicated on graph.

We next comprehensively investigated the cleaved tRNA fragments by deep sequencing (i.e., tRNA reduction and cleavage sequence or TRAC-seq^10,47^) (Fig. 4d, Supplementary Fig. 4b). TRAC-seq identified 23 tRNAs in testes and 22 tRNAs in ovaries that were m^7^G-modified at the RAGGU motif sequence located in the variable loop (Fig. 4e, f, g, Supplementary Fig. 4c). Consistent with the northern blot data (Fig. 4b, c), the TRAC-seq cleavage score was markedly decreased in Mettl1-KO samples, further supporting the function of Mettl1 in m^7^G modification of tRNA (Fig. 4h). Together, these data indicate that Mettl1/Wh is responsible for introducing the m^7^G modification onto a subset of tRNAs in testes.

### *Drosophila* Mettl1 regulates expression of a subset of tRNAs

In *S. cerevisiae*, loss of m^7^G modification on tRNAs results in decreased tRNA abundance^17,22^. This mechanism has been known as the rapid tRNA decay (RTD) pathway mediated by exonucleases^22–24^. Thus, we reasoned that the similar RTD pathway is sparked in *Drosophila* upon the loss of Mettl1. In testes, Mettl1 depletion resulted in changes in the levels of some tRNAs (Fig. 5a, b). In *Drosophila* testis, the mean expression levels of 15 m^7^G-modified tRNAs (ProAGG, ProCGG, ProTGG, ValAAC, ValCAC, ValTAC, LysCTT, LysTTT, ArgTCT, AlaAGC, AlaCGC, AlaTGC, CysGCA, TyrGTA and iMetCAT) were significantly decreased, whereas non-m^7^G modified tRNAs (22 of 45 tRNAs) showed limited changes in the abundance (Fig. 5a). We divided m^7^G-modified tRNAs into two groups according to steady-state levels of tRNA abundance in Mettl1-KO testes (Fig.5b, c, Supplementary Fig. 5a); destabilized in Mettl1-KO (Group 1) and unaltered in Mettl1-KO (Group 2). We also defined m^7^G-hypomodified tRNAs as Group 3 (Fig.5a, c, Supplementary Fig. 5a). Consistent with the TRAC-seq data (Fig. 5a-c), northern blot analysis validated the reduced expression levels of group 1 tRNAs (ProCGG and ValCAC) and the absence of change in no group 2 tRNA (TrpCCA) and group 3 tRNA (LeuAAG) in Mettl1-KO testes (Fig. 5d). (see Figure 7 for the further characterization of these tRNA groups in translation)

**Figure 5.**
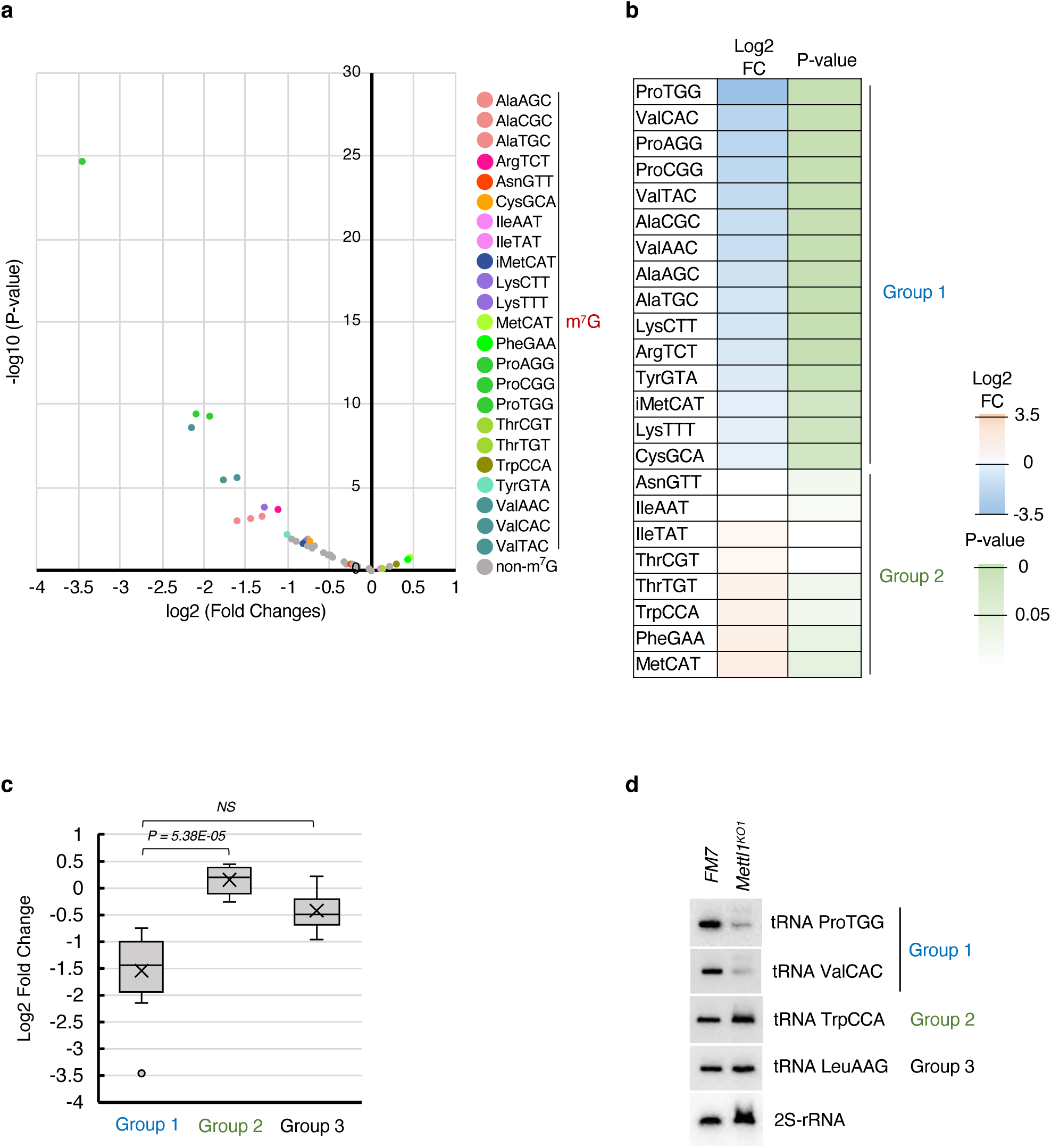
*Drosophila* Mettl1 stabilizes tRNA expression in testis. **(a)** Volcano plot representing changes in tRNA abundance between Mettl1-KO and WT testis. Each dot shows the change of one tRNA abundance. **(b-d)** This study classified *Drosophila* tRNAs into three groups according to difference in abundance between Mettl1-KO and WT, and P-values about significance of difference, Group1 (significantly decreased abundance, log2FC < 0, P-value < 0.05), Group 2 (other m^7^G-modified tRNAs, P-value ≥ 0.05), Group 3 (non-m^7^G tRNA). **(b)** Heatmap showing changes in m^7^G-modified tRNA abundance between Mettl1-KO and WT testis. **(c)** Quantitative comparison of changes in tRNA abundance between three groups of tRNAs defined in this study. Significant differences calculated by Mann-Whitney’s U-test were indicated on graph. **(d)** Northern blot comparison of tRNA expression between Mettl1-KO and WT testes. 2S rRNA is a loading control.

In mammalian A549 cells, *let-7e-5p* miRNA biogenesis is affected by m^7^G methylation^12^, since m^7^G modification in the primary transcript of the *let-7e* disrupts G-quadruplexes formation and facilitates precursor miRNA processing^12^. Since, in *D. melanogaster*, depletion of *let-7* causes a spermatogenesis defect^48^, the phenotype that we observed in Mettl1-KO may be explained by the defect of *let-7* biogenesis in theory. However, the steady-state level of *let-7* miRNA was not significantly changed in *Mettl1*-KO testes, indicating that the sterile phenotype of *Mettl1*-KO males is not caused by m^7^G modification of the *let-7* primary miRNA (Supplementary Fig. 5b). Notably, sequence comparison between mouse and *D. melanogaster* indicates that the mouse RAGGU motif in the *let-7e* primary miRNA in mouse is not identical with that in *D. melanogaster* (UAGGU) and that the G bases that make up the G-quadruplex are A bases in *D. melanogaster*, which might explain why *let-7* was not m^7^G-modified in *Drosophila* (Supplementary Fig. 5c). Taken these observations together, we conclude that the *Drosophila let-7* primary miRNA is not modified by Mettl1 and is unrelated to the sterile phenotype of Mettl1-KO males (Fig. 2, Supplementary Fig. 5b).

### Ectopic expression of eEF1α1 rescues Mettl1-KO infertility

The reduction of a subset of tRNAs in Mettl1-KO led us to investigate whether the RTD pathway explains the sterile phenotype. A genetic approach revealed that Trm8, a METTL1 ortholog in *S. cerevisiae*, is important for m^7^G tRNA modification and stability and for temperature sensitive growth^18,22^. It is noteworthy that the RTD pathway can be suppressed by overexpression of eukaryotic translational elongation factor 1 A (eEF1A) encoded in TEF1 gene in yeast because eEF1A directly bind to tRNAs and protects them from RNA decay machineries^49,50^. Therefore, we asked whether overexpression of eukaryotic translational elongation factor 1 alpha 1 (eEF1α1, CG8280), a fly homolog of eEF1A, rescues the male sterility in Mettl1-KO flies (Fig. 6a). As expected, the overexpression of eEF1α1 significantly recovered the fertility of the Mettl1-KO males, indicating that suppression of the RTD pathway is critical for *Drosophila* fertility (Fig. 6a, b).

**Figure 6.**
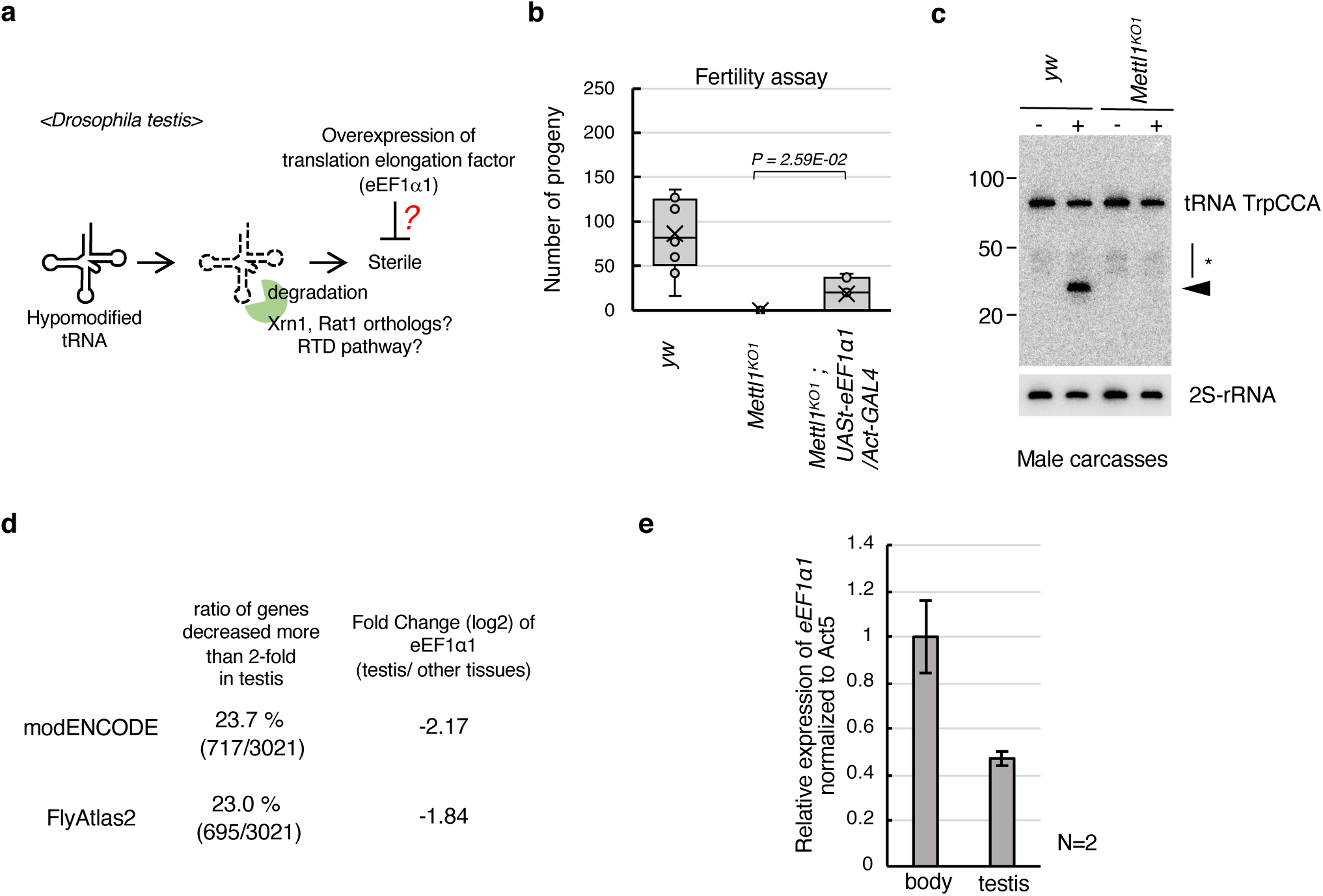
Ectopic expression of eEF1a recovered sterility of Mettl1-KO. **(a)** Hypothesized model of the relationship between eEF1α1, the RTD pathway and m7G target tRNAs determining fertility of *D. melanogaster*. **(b)** Number of progenies gained from male fertility assay using WT (*yw*), Mettl1-KO (*Mettl1^KO1^*), Mettl1-KO with overexpression of eEF1α1 (*Mettl1^KO1^; UASt-eEF1α1*/*Act-Gal4*). Significant differences calculated by two-tailed Student’s t-test were indicated on graph. (**c)** Northern blot validation of m^7^G modification in tRNA TrpCCA from male carcasses sample. (**d)** Ratio

To explore the function of Mettl1 in nongonadal tissues, we performed western blot and northern blot analyses of male carcasses from which testes were manually removed. These analyses showed that Mettl1 and m^7^G-modified tRNAs were present in nongonadal tissues (Fig. 6c, Supplementary Fig. 2d). Notably, the only defect that was apparent in Mettl1-KO flies was in spermatogenesis (Fig. 2). To explore potential reasons why defects are restricted to gonad function, we carefully studied the mRNA levels of eEF1α1. From the tissue-specific gene expression profile of FlyBase, we reanalyzed and found that eEF1α1 showed lower expression levels in testis (log2FC = −2.17 from RNA-seq dataset in modENCODE; log2FC = −1.84 from that in FlyAtlas2) (Fig. 6d, Supplementary Fig. 6). Consistent with this result, RT-qPCR data showed a lower eEF1α1 expression in testes than that in other tissues (Fig. 6e). Given that eEF1α1 may compete with the RTD pathway, a lower level of eEF1α1 mRNA indicates that the testes may be more susceptible than other tissues to RTD pathway-dependent degradation.

### Translatome analysis in Mettl1-KO testis

To examine whether the reduction of tRNAs affects mRNA translation, we performed a ribosome profiling from dissected testes^51,52^. Given the limited materials of dissected testes, we employed a method tailored for low input with RNA-dependent RNA amplification^53^. Remarkable 3-nt periodicity (Supplementary Fig. 7a) and high reproducibility (Supplementary Fig. 7b, c) ensured the high quality of data even in the tiny fly tissues. We observed the remarkably high ribosome occupancy on A-site codons by the loss of Mettl1 (Fig. 7a). Many of these codons corresponded to codons decoded by the m^7^G-modified tRNAs (Fig. 7a). Notably, among the m^7^G-modified tRNAs, the sub-group subjected to RTD (group 1, Fig. 5c) showed the prominent effects on ribosome traversal (Fig. 7b). Strikingly, ribosome pausing on CCA codon, recognized by group 1 tRNA ProTGG, induced the collisions of trailing ribosomes, forming disomes and trisomes (Fig. 7c). We observed that more pronounced ribosome stalling on CCA, GUG, CCU, and CCG codons occurred on a subset of transcripts such as ones encoding genes essential for fertility (Fig. 7d, Supplementary Table 1). In addition to the low expression of eEF1α1, this may also strengthen phenotype in testes by the loss of Mettl1.

**Figure 7.**
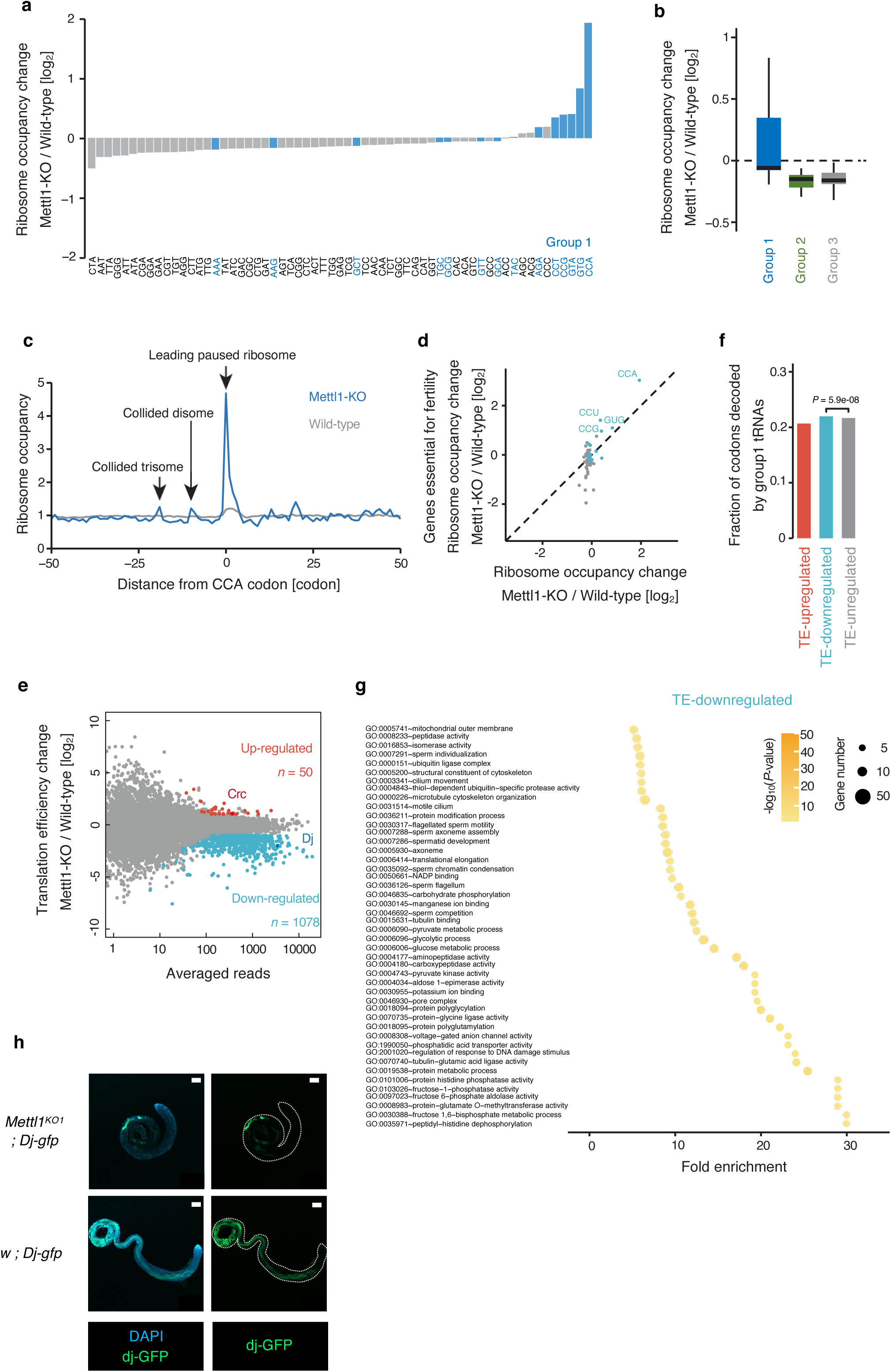
Mettl1 maintains proper translatome to preserve complete fertility in gonads. **(a)** Ribosomal occupancy change across A-site codons by Mettl1-KO in testis. **(b)** Ribosome occupancy changes in the codons decoded by indicated tRNA groups. **(c)** Metagene plot for ribosome occupancy around CCA codons. The positions of paused and collided ribosome are highlighted by arrows. **(d)** The difference of ribosome occupancy change on A-site codons by Mettl1-KO between all analyzed genes and genes that have been reported to be essential for fertility. **(e)** MA (M, log ratio; A, mean average) plot showing the fold change of translation efficiency by Mettl1-KO and the averaged read number. Each dot represents each transcript. Significantly changed transcripts (upregulate, log_2_ fold change ≥ 1 and FDR ≤ 0.05; down-regulated, log2 fold change ≤ −1 and FDR ≤ 0.05) are highlighted. **(f)** Fraction of codons decoded by group 1 tRNAs in the indicated transcript group. The significance was calculated by Pearson’s χ^2^ test. **(g)** Gene ontology associated with TE down-regulated genes by Mettl1-KO (p-value < 0.01). **(f)** Immunostaining comparison of Dj-GFP expression between control (*w ; Dj-GFP*) and Mettl1-KO (*Mettl1^KO1^; Dj-GFP*) testes.

In addition to the codon-wise assessment, we also investigated the impact of m^7^G tRNA modification on translation efficiency across ORFs. Normalizing the ribosome footprints by RNA abundance measured by RNA-Seq (Supplementary Fig. 7d), we measured translation efficiency (TE) and that those in a notable number of transcripts (1078) were down-regulated by the loss of METTL1 (Fig. 7e). These TE down-regulated transcripts were more prone to possess codons decoded by group 1 tRNAs (Fig. 7f). Gene ontology (GO) associated with the TE down-regulated genes included the key terms for gonadal functions, such as sperm individualization, sperm axoneme assembly and spermatid development (Fig. 7g), consistent with the observed phenotype of Mettl1-KO. We noticed translational impairment of Don juan (Dj), which has been reported to be expressed in elongated spermatids and individualized sperms^54^, in Mettl1-KO (Fig. 7e). We validated this observation by the transgene of GFP-tagged Dj; the Dj-GFP expression significantly decreased in Mettl1-KO testis (Fig. 7h).

In addition to TE down-regulated transcripts, we found a small fraction of transcripts with higher TE by Mettl1-KO (Fig. 7e). These mRNAs included Crc, a *Drosophila* homolog of ATF4, which bears upstream ORF (uORF) (Supplementary Fig. 7d)^55^. Although the uORF typically traps scanning ribosomes and inhibits the complex reaching to downstream main ORF, the reduction of initiator tRNA (Fig. 5b) may explain the induction of leaky scanning of uORF as represented by eIF2 inactivation occurred in integrated stress response^56^. Taken together, these data support the translational changes at the late stage of spermatogenesis leads to the infertility of Mettl1-KO male (Fig. 1d).

## Discussion

Here, we addressed the essential role of Mettl1 in the formation of elongated spermatids and male fertility through m^7^G tRNA modification. We found that ribosome pausing occurs at Pro and Val codons decoded by m^7^G-modified tRNAs in Mettl1-KO testes. Consistent with defective elongated spermatid formation in Mettl1-KO testes, Mettl1 might play an important role in the mRNA translation of mitochondrial derivatives involved in sperm tail formation. During late-stage spermatogenesis, chromatin modifiers cause chromosome compaction to shut down transcription. This stage therefore relies on translation of stored mRNAs, which is regulated by RNA binding proteins^57,58^. Our data show that epitranscriptomic regulation of tRNA, m^7^G modification, also plays a critical role during this process. Furthermore, no apparent developmental defect, apart from to fertility, was observed despite the presence of Mettl1 and m^7^G modified tRNAs in nongonadal tissues. Notably, we showed that eEF1α1, an inhibitor of the RTD pathway, is expressed at low levels in testes, and its overexpression restored fertility to Mettl1-KO males. This indicates that the balance between the activity of the RTD pathway and the expression levels of RTD pathway competitors, such as eEF1α1, determines the steady-state level of hypomodified tRNAs and the tissues where m^7^G-tRNA modifications are particularly critical for normal development.

The finding that Mettl1-KO resulted in decreased expression of the majority, but not all, m^7^G-modified tRNAs (Fig. 5a, b) was consistent with previous studies in mammals and yeast^10,22,27^. Critical components of the RTD pathway in yeast, Rat1/Dhp1, Met22, and Xrn1 are all present in higher eukaryotes^59^. Therefore, the molecular machinery reported in the yeast RTD pathway may also function in other organisms, although this has not been addressed in animals. We showed that eEF1α1 overexpression rescues Mettl1-KO fertility, which supports a similar system working in animals. Interestingly, our data show the m^7^G-modified tRNA species to be largely similar between *Drosophila* and mammals, while there were some differences in Mettl1 dependency for the effects on stability among tRNA species. For example, m^7^G-modified tRNA PheGAA was stable in *Drosophila* Mettl1-KO (Fig. 5a, b) but reduced in METTL1-depleted mammalian cells^10,27^. RTD acts on tRNAs lacking one or more of several different modifications^49^; therefore, we speculate that these differences are caused by varying combinations of other tRNA modifications. Indeed, a subset of hypomodified tRNAs is reduced in *Drosophila* adult flies lacking cytosine-5 tRNA methyltransferases^60^; therefore, it is likely that tRNA decay can occur in diverse cell types, depending upon the combinations of RNA modifications.

It was surprising to find that the only obvious abnormality in Mettl1-KO flies was restricted to fertility because Mettl1 is ubiquitously expressed and m^7^G modification of tRNAs also occurs in nongonadal tissues (Fig. 6c). As discussed above, RTD pathway proteins are conserved in *Drosophila* and are ubiquitously expressed. We suggest that unmodified testicular tRNAs are more susceptible to degradation compared with those in other tissues because ectopic expression of eEF1α1, a competitor of the RTD pathway, restored the phenotype (Fig. 6b). Notably, there is no post-meiotic transcription in *Drosophila* testes, which is conserved in mammals^61^. Therefore, proteins involved in spermatid differentiation must be translated from previously produced mRNAs that are stored in a translationally repressed state. During this process, numerous mRNAs are translated and spermatids undergo dynamic morphological changes, which include nuclear condensation and axoneme formation, without supplementation of fundamental translation factors, such as translation elongation factors. Based on our results showing that elongated spermatid formation is impaired in Mettl1-KO males and that eEF1α1 overexpression restored fertility, it is likely that the low eEF1α1 expression may sensitize testes than other tissues to Mettl1 deficiency (Fig. 6e). However, we cannot rule out the possibility that the RTD pathway is stronger in testes than in other tissues and that the expression levels of the tRNAs for each codon differ between testis and other tissues. It is therefore necessary to understand the molecular details of the RTD pathway and the interplay with its competitors at the tissue and single cell level. Notably, it remains controversial whether METTL1 depletion affects global translation efficiency. METTL1 depletion in human glioblastoma LNZ308 cells^27^, HuCCT1 and RBE cells^26^, and in mouse ES cells^10^, reduces global translation efficiency. In contrast, METTL1-depletion did not affect overall translation levels in A549, HepG2 and HeLa cells^12,15,28^. Considering that initiator tRNA was classified into RTD target in our analysis, global reduction of translation might be explained by the resultant reduction of initiation rate. Alternatively, global translation shut-off caused by ribosome stalling (i.e., integrated stress response, ISR)^62^ may be sparked. It is possible that these processes induced by METTL1 depletion may depends on cell types. Given that leaky scanning in the mutant and eEF1α1-mediated phenotype rescue, both processes could be induced in the fly testes.

In addition to the potential global effect, we observed that a subset of mRNA’s translation was selectively affected in TEs. Given that translation initiation speed is a general rate limiting step of protein synthesis and defines the overall footprint counts from ORFs^63^, the reduction of TE should stem from the decrease of translation initiation rate. The correspondence between ribosome pausing and reduced translation initiation rate may be explained by the feedback mechanism that senses ribosome collision^64,65^. The cellular component term analysis of genes that were down-regulated in Mettl1-KO testes revealed that Mettl1 is critical for translation of mRNAs involved in elongated spermatid and mature sperm formation, which is supported by our experimental observations (Fig. 2). The biological process term analysis of genes that were down-regulated in Mettl1-KO testes revealed that Mettl1 is critical for the translation of mRNAs involved in the catabolism of fructose, which acts as an energy source for spermatozoa^66^. These data indicate that Mettl1-KO severely impairs protein synthesis from a subset of mRNA during late rather than early stages of spermatogenesis, resulting in male sterility.

Interestingly, our ribosome profiling analyses showed increased translation efficiency of 50 transcripts (Fig. 7e). A recent report revealed that internal m^7^G-modified mRNAs with GA-rich motifs are directed to stress granules via Quaking proteins in human cells^28^. Stress granules contain nontranslating mRNAs^67^; therefore, the high TE in Mettl1-KO testes might result from depression of stress granule-stored, m^7^G-modified mRNAs.

Activation of Mettl1 can occur in cancer cells, resulting in stabilization of tRNA (ArgTCT4-1) and increased translation of cell-cycle gene mRNAs, such as CDK6 and others, which promotes tumor progression^26,27^. Notably, not only METTL1/WDR4 but also eEF1α1 expression is elevated in hepatocellular carcinoma cell lines and clinical samples^68^, indicating that tRNA stability (ArgTCT4-1) might be synergistically increased in hepatocellular carcinoma cells. Tumor progression depends on the promotion of tRNA (ArgTCT4-1) methylation by METTL1; therefore, either reduction of eEF1α or activation of the RTD pathway might attenuate tumor progression. A combination of some drugs repressing the RTD pathway machinery may enhance the effects of Mettl1-inhibitor.

## Materials and Methods

### Fly stocks

A complete list of fly stocks used in this study is presented in Supplementary Table 2. All stocks were maintained at 25°C on standard medium. Mettl1-knockout (Mettl1-KO) lines were generated using the transgenic CRISPR-Cas9 method^69^. To generate mutants, we used the following strains: *y^1^ v^1^ nos-phiC31; attP40* host (NIG-FLY stock TBX-0002), *y^2^ cho^2^ v^1^*; *Sp hs-hid/CyO* (NIG-FLY stock TBX-0009), *y^1^ w^1118^; +; attP2{nos-cas9}*^70^, and *Df(1)JA27/FM7c, P{w[+mC]=GAL4-Kr.C}DC1, P{w[+mC]=UAS-GFP S65T}DC5, sn[+]* (Bloomington Drosophila Stock Center #5193). Disruption of the Mettl1 region in mutants was validated by PCR and Sanger sequencing. All crosses in this study were performed at 25°C. Oligonucleotide sequences for Mettl1-KO line generation are shown in Supplementary Table 3. To confirm the requirement of *Wh* in Mettl1-dependent m^7^G methylation, we used the following strain: *w* wuho*^56^*/FM7c, sn^+^* (Bloomington Drosophila Stock Center #41112). *w* / Y ; P{Dj-GFP.S}AS1 / CyO* (Bloomington Drosophila Stock Center #5417) was used to validate expression of *Dj* (*don juan*) in testes.

### Fertility assay

To test male fertility, a single male fly was mated with three control (*y^1^ w^1118^*) females at 25°C for three days. To test female fertility, a single female was mated with three control (*y^1^ w^1118^*) males at 25°C for 10 days. After mating, the parental flies were removed, and incubation continued for 11 days in the male fertility assay, and for 4 days in the female fertility assay. Progeny were counted and the average number per vial calculated. Ten independent vials for each strain were prepared. The complete list of flies used in these assays is shown in Supplementary Table 2.

### Western blotting

Western blotting was performed as described previously^71^. Ovaries were collected from 3–4-day-old adult females. An anti-Mettl1 monoclonal antibody (generated in this study), an anti-Wh monoclonal antibody (generated in this study), and an anti-Tubulin antibody (E7 culture supernatant, Developmental Studies Hybridoma Bank [DSHB]) were used as primary antibodies. A goat anti-mouse IgG (H+L) Poly-HRP secondary antibody (Thermo Fischer, USA, #32230, 1:5000–10000) was used the secondary antibody.

### Measurement of ovary size

Ovaries were dissected from 7-day-old adult females. Ovaries were fixed in 4% paraformaldehyde phosphate buffer solution (Nacalai Tesque, Japan) for 15 min at room temperature or overnight at 4°C. After washing with PBS-T three times, ovary size was measured using a stereomicroscope and cellSens (Olympus, Japan) imaging software. Significant differences in ovary size were evaluated by two-tailed Student’s t-test.

### Immunofluorescence

Immunostaining analysis was conducted as described previously ^72^. Testes from 3-day-old adult males and ovaries from 7-day-old adult females were dissected in ice-cold PBS. Tissues were fixed in 4% paraformaldehyde phosphate buffer solution (Nacalai Tesque #09154-85) for 15 min at room temperature. The fixed tissues were washed five times with PBS containing 0.1% Tween 20 (PBS-T), and then incubated two times in PBS-T containing 0.2% Triton X-100 and 0.1% BSA (PBS-BT) for 10 min. The washed tissues were then incubated with primary antibodies in PBS-BT at 4°C for 1 day. Primary antibodies used were: anti-Aub (4D10) ^73^, anti-acetylated tubulin (Sigma, 1:100 dilution), anti-boule (1:250 dilution, a kind gift from S. Wasserman, University of California San Diego), anti-FLAG (M2, Sigma, 1:1000 dilution; to detect Mettl1-3×Flag), anti-Fas3 (7G10, DSHB), and anti-Tj (10009Ab-1, NIG-FLY, Japan). Incubated tissues then were washed five times and further washed two times with PBS-BT. Next, the tissues were incubated at 4°C for 1 day with the following secondary antibodies as appropriate: Alexa 546-conjugated anti-mouse IgG (Thermo Fisher, A11030, 1:250–1000 dilution), Alexa 488-conjugated anti-mouse IgG (Thermo Fisher, A11029, 1:250 dilution), and Alexa 546-conjugated anti-rabbit IgG (Thermo Fisher, A11071, 1:250–1000 dilution). After washing three times for 10 min in PBS-T, tissues were mounted in Vectashield Mounting Medium containing DAPI (Vector Laboratories). Images of testes and ovaries were taken using a confocal microscope (Zeiss LSM900). Seminal vesicles and elongated spermatids in testes were fixed by the same method and observed by light field confocal microscopy (Zeiss LSM900).

### Production of monoclonal antibodies

Recombinant full-length *Drosophila* Mettl1 protein tagged with maltose binding protein and Wh (N-terminal 200 amino acids) tagged with glutathione S-transferase (GST) were prepared as antigens and purified from *E. coli*. Mice were immunized with each antigen. The production and selection of hybridomas were performed as previously described^74^.

### Plasmid construct

To produce plasmids expressing *Drosophila* Mettl1, Mettl1-Catalytic dead (Cd), Mettl1-3×Flag, nosp-Mettl1, UASt-Mettl1 and UASt-eEF1α1, Mettl1 and eEF1α1 coding regions were amplified using KOD plus DNA polymerase (Toyobo #KOD-201). The Nanos promoter region was amplified from the following vector: pBFv-nosp-Cas9. The amplified fragments and vector pBFv-UAS3 were fused using the In-Fusion® HD Cloning Kit (TaKaRa). The oligonucleotide sequences used are shown in Supplementary Table 2. To produce the plasmids expressing Mettl1 and Wh in *E. coli*, the Mettl1 and Wh coding sequences were amplified using KOD plus DNA polymerase (Toyobo) and were then integrated into pGEX 5x-1 (Amersham), or pMAL vector (NEB). The oligonucleotide sequences used are shown in Supplementary Table 3.

### GST pull-down assays

GST pull-down assays were performed as previously described^75^. Purified GST- or MBP-tagged proteins were incubated in methylation buffer (20 mM Tris-HCl pH 8.0, 100 mM NaCl, 0.2 mM DTT, 10 mM EDTA, 0.2 mM MgCl_2_) with Glutathione Sepharose™ 4B (GE healthcare 17-0756-01) or Amylose Resin (NEB, E8021S) for 1 h at 4°C to immobilize GST- and MBP-proteins, respectively. Immobilized proteins were washed three times with methylation buffer. After washing, protein to be tested was added and incubated for 4h at 4°C. After incubation, proteins were washed five times with methylation buffer, and then eluted with 2×Sample buffer [100 mM Tris-HCl (pH 6.8), 4% (w/v) SDS, 0.2% (w/v) bromophenol blue, 20% (v/v) glycerol, and 200 mM DTT]. Samples were incubated for 5 min at 95°C, and were then resolved by sodium dodecyl sulfate-polyacrylamide gel electrophoresis (SDS-PAGE). The resolved proteins were detected by Coomassie brilliant blue staining.

### Preparation of protein samples for LC-MS/MS analysis

For the preparation of immunoprecipitated samples for LC–MS/MS, the anti-3×FLAG antibody used for immunoprecipitation was cross-linked to beads by dimethyl pimelimidate (Thermo Fisher). Immunoprecipitation was performed using mouse anti-FLAG M2 or mouse non-immune IgG (negative control) and ovaries from negative control (Oregon R) or 3×Flag tagged Mettl1(*attp40[Mettl1-Flag]*). We used two different buffer conditions: Tris-Triton buffer [20 mM Tris-HCl (pH 8.0), 100 mM KCl, 5 mM MgCl_2_, 2 mM DTT, 0.1% TritonX-100] or Hepes-NP40 buffer [30 mM Hepes-KOH (pH 7.3), 150 mM KOAc, 5 mM MgOAc, 5 mM DTT, 0.1% NP40]. LC-MS/MS analysis was performed as following previous paper^76^. The immunoprecipitated proteins were run on 4-12 % SDS-PAGE gel by 1 cm from the well and stained with SimplyBlue (Thermo Fisher) for in-gel digestion. The gel containing proteins was excised, cut into approximately 1 mm sized pieces. Proteins in the gel pieces were reduced with DTT (Thermo Fisher), alkylated with iodoacetamide (Thermo Fisher), and digested with trypsin and Lysyl endopeptidase (Promega) in a buffer containing 40 mM ammonium bicarbonate, pH 8.0, overnight at 37°C. The resultant peptides were analyzed on an Advance UHPLC system (AMR/Michrom Bioscience) connected to a Q Exactive mass spectrometer (Thermo Fisher) processing the raw mass spectrum using Xcalibur (Thermo Fisher Scientific). The raw LC-MS/MS data was analyzed against the UniprotKB database restricted to *Drosophila melanogaster* using Proteome Discoverer version 1.4 (Thermo Fisher) with the Mascot search engine version 2.5 (Matrix Science). A decoy database comprised of either randomized or reversed sequences in the target database was used for false discovery rate (FDR) estimation, and Percolator algorithm was used to evaluate false positives. Search results were filtered against 1% global FDR for high confidence level. Quantitative analyses were performed using scaffold 5 (Proteome Software Inc, Portland, OR), and volcano plots of identified proteins were generated using student’s t-test or fisher’s exact test under a normalization scheme.

### Northern blotting

Northern blotting was performed as described previously^77^. Sample RNAs were denatured with NorthernMax Formaldehyde Load Dye (Invitrogen #AM8552) for 3 min at 65°C. Sample RNAs were resolved by electrophoresis on 12% urea-PAGE gels, which contain 7 M urea, 10× TBE (1 M Tris base, 1 M boric acid, 0.02 M EDTA), and 12% acrylamide. To assess m^7^G site-specific cleavage of tRNAs and to confirm tRNA levels, 1.25 μg (from ovaries) and 0.5 μg (from testes) sample RNAs were resolved by electrophoresis and then transferred to Hybond N+ membranes (Cytiva). Transferred RNAs were UV-crosslinked with 1200×100 mJ/cm2 irradiance at 254 nm. After crosslinking, hybridization was conducted at 42°C overnight in 7% SDS, 0.2 M sodium phosphate (pH 7.2), and 1 mM EDTA with ^32^P-labeled DNA probes. Membranes were then washed with 2× saline sodium citrate containing 0.1% SDS at 42°C. To detect *in vitro* m^7^G methylation, the hybridization step was skipped. The oligonucleotide sequences used are shown in Supplementary Table 3.

### In vitro methylation assay

GST, GST-Wh, MBP, MBP-Mettl1 and MBP-Mettl1-Cd were produced and purified from *E. coli*. *In vitro* tRNA methylation assay was performed essentially as described previously with some modifications^44,75,78^. Purified GST and MBP protein (1 μg each) were incubated with chemically synthesized RNA (0.3 nmol) at 26°C in a buffer containing 50 mM Tris-HCl (pH 8.0), 100 mM KCl, 5mM MgCl_2_, 2 mM DTT and 2 μCi/mL S-adenosyl-L-[methyl-^14^C] methionine (PerkinElmer). After 3h incubation, RNAs were isolated with phenol:chloroform and ethanol-precipitated. The resultant RNAs were run on 7 M urea containing denaturing polyacrylamide gels. The gels were stained with Toluidine Blue *O* in 0.5 X TBE, destained with 0.5 X TBE, and dried. The ^14^C-labeled RNA bands are visualized on a Typhoon FLA900 (GE Healthcare).

### m7G site-specific reduction and cleavage

Total RNA from ovaries and testes were isolated using Isogen (Nippon Gene, 311-02501) following the manufacturer’s instructions. Ten micrograms of total RNA were incubated with 0.1 M NaBH_4_ and 1 mM free m^7^GTPfor 30 min on ice in the dark. Reduced RNAs were precipitated with 3 M sodium acetate pH 5.2 (Thermo Fisher), glycogen (Nacalai Tesque) and ethanol at −20°C for at least 1 hr. RNA samples reduced by NaBH_4_ were then reacted with aniline-acetate solution (H_2_O:glacial acetate acid:aniline, 7:3:1) at room temperature in the dark for 2 h to cause m^7^G site-specific cleavage^45,46^. The cleaved RNAs were detected by northern blotting. Total RNA from Wh-KO males was isolated from 3-day-old adult male testes of *w* wh^56^/FM7c sn+* flies (Bloomington Drosophila Stock Center #5417).

### TRAC-seq and profiling of tRNA expression

Sequencing and data analysis were performed as previously described^47^ with some modifications. Small RNAs (< 200 nt) were isolated from whole ovary (one sample) and testis (three replicates) RNAs using a mirVana miRNA Isolation Kit (Thermo Fisher, AM1561) following the manufacturer’s instructions. To exclude methylation inhibiting the reverse transcription of tRNAs, 2.5 μg of extracted small RNAs were treated with recombinant AlKB and AlkB D135S proteins. Demethylated small RNAs were reduced and cleaved by the method described in the “m^7^G site-specific reduction and cleavage” section of the manufacturer’s protocol. After cleavage and ethanol precipitation, small RNA sequencing libraries of m^7^G tRNAs were constructed using a NEBNext Multiplex

Small RNA Library Prep Set for Illumina (NEB, E7300S). In preparing the library, TGI-RT (Thermostable Group II intron reverse transcriptase, TGIRT-III-10, InGex) was used instead of the reverse transcriptase in the library kit. The prepared libraries were sequenced on a Novaseq 6000 system. The 290 reference sequences of mature *Drosophila* tRNAs (*i.e.* dm6-mature-tRNAs) were downloaded from GtRNAdb^79^. Identical sequences were removed to yield 84 unique sequences and a “CCA” sequence was added to the 3′-end of each tRNA (*i.e.* “dm6-mature-tRNAs_unique_CCA”). The 5′- and 3′- adapter sequences of the sequenced reads were trimmed and low-quality sequences were filtered using FASTP ver. 0.23.2^80^ to obtain the “clean reads”. The clean reads were mapped to the reference “dm6-mature-tRNAs_unique_CCA” using BOWTIE ver. 1.3.1^81^. From the mapping data (bam), the number of reads mapped on each tRNA (*i.e.* “TotalReads”) was obtained using PICARD ver. 3.0.0 (https://broadinstitute.github.io/picard/command-line-overview.html). The number of reads starting at each site of each tRNA (*i.e.* “StartReads”) was obtained using the custom shell script. The ratio between StartReads and TotalReads as an indicator of the m^7^G-dependent read cleavage (i.e. “StartRatio”) was calculated for each site *i* of each tRNA of chemically treated and non-treated samples as:

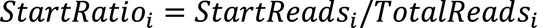

The cleavage score as an overall indicator of the m^7^G modification (*i.e.* “CleavageScore”) was calculated for each site *i* of each tRNA of WT and Mettl1-KO samples as:

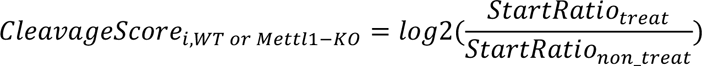

The difference in cleavage score between WT and Mettl1-KO samples (*i.e.* “Cleavage score”) was defined as another indicator of Mettl1-dependent m^7^G modification as:

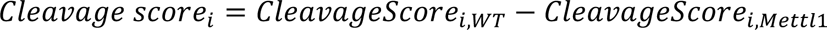

For testes, these feature values were averaged from the data of three replicates. The sites with StartRatio_WT,treat_ ≥ 0.02 were regarded as candidates. Those with CleavageScore_i, WT_ ≥ 2 and Cleavage score_i_ ≥ 2 were extracted as the final candidate m^7^G-dependent cleavage sites. The sequences of ± 10 bases around the candidate m^7^G site were analyzed to detect the conserved motif for the m^7^G modification using MEME ver. 5.5.2^82^.

The chemically non-treated testis samples for TRAC-seq (WT: three replicates; Mettl1-KO: three replicates) were processed to profile tRNA expression using the tRAX pipeline ^83^.

### Reanalysis of tissue-specific gene expression profiles in FlyBase

The two datasets (modENCODE Anatomy RNA-Seq dataset: FBlc0000206 and FlyAtlas2 Anatomy RNA-Seq dataset: FBlc0003498) were downloaded from FlyBase (http://ftp.flybase.org/releases/FB2022_04/precomputed_files/genes/gene_rpkm_report_fb_2022_04.tsv.gz) to search for “ubiquitously expressed and low variance (ULV)” genes. These data include the reads per kilobase per million mapped reads (RPKM) of each gene for each tissue sample. In one approach, the RPKM data were directly used for the subsequent analysis. In another approach, the RPKM values were TMM-normalized using the EDGER package^84^ as is performed for identifying housekeeping genes in human and mouse^85^. The ULV genes were defined using the following moderate criteria: 1) The genes are expressed in all tissues (*i.e.* non-zero value). 2) The genes have a low variance of expression (standard deviation of the log2 RPKM < 1.3) across all tissues except testes. The fold change of expression of each ULV gene in the testis was calculated against the mean of all tissues except the testis. The genes with a log2 fold change < –1 in the testis were regarded as “significantly decreased” compared with the mean expression level.

### Ribosome footprinting and RNA-Seq

The cell lysate was prepared as previously reported with modifications^86^. Testes were dissected from 2-3-day-adult-males in ice-cold PBS with 100 µg/ml cycloheximide. Dissected testes were immediately frozen with liquid nitrogen and stored at −80°C. We prepared 600 pairs of testes per sample and two replicates from wild-type (*yw*) and Mettl1-KO (*Mettl1^KO1^*) males, respectively. Frozen testes were mixed with 300 µl of frozen droplets of lysis buffer (20 mM Tris-HCl pH7.5, 150 mM NaCl, 5 mM MgCl_2_, 1 mM dithiothreitol, 1 % Triton X-100, 100 µg/ml chloramphenicol, and 100 µg/ml cycloheximide), and were then ground at 3,000 rpm for 15 s using a Multi-beads Shocker (YASUI KIKAI). The lysate was slowly thawed at 4°C and treated with 10 U of Turbo DNase (Thermo Fisher Scientific) on ice for 10 min to remove the genome DNA. The supernatant was further centrifuged at 20,000 g for 30 min at 4°C.

Library construction of ribosome profiling (or Thor-Ribo-Seq) was conducted as following previous protocols^53,86^. We treated cell lysate containing 8.5 µg total RNA with 20 U RNase I (Epicentre) in a 300 µl reaction at 25°C for 45 min. Then, the reaction was stopped by 200 U of SUPERase•In RNase Inhibitor (Thermo Fisher Scientific). Ribosomes were isolated by sucrose cushion ultracentrifugation at 100,000 rpm at 4°C for 1 h using TLA110 rotor (Beckman Coulter). RNAs in the ribosome pellet were isolated with TRIzol-LS (Thermo Fisher Scientific) andDirect-zol RNA Microprep kit (Zymo Research). Purified RNA fragments ranging 17 to 34 nt were selected by gel electrophoresis, dephosphorylated by T4 polynucleotide kinase (New England Biolabs), ligated to preadenylated linkers containing the T7 promoter region by T4 RNA ligase 2, truncated KQ (New England Biolabs). Before ligation, the linkers were preadenylated by Mth RNA Ligase (NEW England Biolabs). Linker-ligated RNA were treated with riboPOOL for *D. melanogaster* (siTOOLs Biotech) to remove the fragments of ribosomal RNAs. Then, we performed *in vitro* transcription using a T7-Scribe Standard RNA IVT Kit (CELLSCRIPT). The RNAs were dephospholylated by T4 polynucleotide kinase (New England Biolabs) and ligated with linker for reverse transcription. The cDNAs were synthesized by ProtoScript II (New England Biolabs) and amplified using Phusion High-Fidelity DNA Polymerase. The DNA libraries were sequenced on HiSeq X (Illumina) with an option of paired-end 150-bp.

For RNA-Seq, we extracted total RNA from the lysate originally used for ribosome profiling using TRIzol LS (Thermo Fisher Scientific) and the Direct-zol RNA MicroPrep Kit (Zymo Research). We then utilized 1 µg of the total RNA for the SEQuoia Express Standard RNA Library Prep Kit (Bio-Rad). rRNA depletion was performed with riboPOOL for *D. melanogaster* (siTOOLs Biotech).

Sequence data were processed using previously established methods^87,88^. Translation efficiency change was determined using the DESeq2 package^89^.

## Acknowledgments

We thank T. Suzuki for critical suggestions on this project; and A. Terui, A. Masuda, Y. Kitahara, Y. Nagashima, and T. Miyoshi for technical assistance; and M. Masukawa, S. Kobayashi, the Drosophila Genetic Resource Center in Kyoto Institute of Technology and the Bloomington stock center for providing the fly strains and Jeremy Allen, PhD, from Edanz (https://jp.edanz.com/ac) for editing a draft of this manuscript. This work was partially supported by JSPS KAKENHI under grant numbers JP18H02379, JP16H06279 (both PAGS), JP23H02415 (SI) and JP22H02669 (KS); and the Takeda Science Foundation, the Yamada Science Foundation, the Mochida Memorial Foundation (KS), The Naito Science & Engineering Foundation and the Uehara Memorial Foundation (KS); AMED under grant number JP23gm1410001 (SI); and the Program of the Joint Usage/Research Center for Developmental Medicine and High Depth Omics, IMEG, Kumamoto University (KS). K.T. was a recipient of RIKEN Student Researcher program and a World-leading Innovative Graduate Study Program in Proactive Environmental Studies (WINGS-PES) from The University of Tokyo. This study used the HOKUSAI SailingShip supercomputer facility at RIKEN.

## Author contributions

**Shunya Kaneko**: Conceptualization; formal analysis; investigation; methodology; visualization; validation, writing-original draft; writing-review and editing. **Keita Miyoshi**: investigation; methodology; visualization; validation, writing-review and editing. **Kotaro Tomuro**: Data curation, investigation; methodology; writing-review and editing. **Makoto Terauchi**: Data curation, software; investigation; methodology; writing-original draft; writing-review and editing. **Shu Kondo**: Resource; methodology and editing. **Naoki Tani**: LC/MS/MS; methodology and editing. **Kei-Ichiro Ishiguro**: Investigation; methodology and editing. **Atsushi Toyoda**: Methodology. **Hideki Noguchi**: Data curation, investigation; methodology; writing-original draft; writing-review and editing. **Shintaro Iwasaki**: Data curation, investigation; methodology; writing-original draft; writing-review and editing. **Kuniaki Saito**: Conceptualization; funding acquisition; investigation; methodology; project administration; supervision; writing-original draft; writing-review and editing.

## Data and code availability

The original raw data of mass analyses in the manuscript are deposited in ProteomeXchange (PXD044501) / jPOST repository (JPST002287). The TRAC-Seq, ribosome profiling, and RNA-Seq data obtained in this study were deposited in the National Center for Biotechnology Information (NCBI) database (GSE241519, ribosome profiling and RNA-Seq) and DNA Data Bank of Japan (DDBJ) (DRR499643-DRR499658, TRAC-seq).

