## Supplementary Information for "Mettl1-dependent m^7^G tRNA modification is essential for maintaining spermatogenesis and fertility in *Drosophila melanogaster*"

**Supplementary Figure S1. RNA methyltransferase-like domain in eukaryotic Mettl1 is highly conserved in *Drosophila* Mettl1 and Mettl1 is important for ovary development.**

**(a)** Amino acid alignment of Mettl1 from *Drosophila* and other eukaryotes, including *Homo sapiens*, *Mus musculus* and *Danio rerio*. Blue underlined region indicates the RNA methyltransferase-like domain. Red box indicates the amino acid residue that is responsible for RNA methyltransferase activity. **(b)** Amino acid sequences of the products expressed from *Drosophila Mettl1*. Mettl1-KO1 expressed a product containing less nonsense sequence than KO2; therefore, we mainly used the KO1 strain as the Mettl1-KO line. Orange and green letters indicate nonsense sequences. **(c)** Western blot showing Mettl1 and Wh expression in *Drosophila* ovaries. **(d)** Schematic of the fertility assay. One test fly and opposite sex WT (*yw*) flies were prepared for each vial. For male fertility assays, vials were incubated for 3 days and parental flies were then removed. The vials were then incubated for a further 11 days. For female fertility assays, vials were incubated for 10 days, and then the parental flies were removed. The vials were then incubated for a further 4 days. **(e)** Representative images of ovaries from control (*$Mettl1^{KO1}$/FM7*, left), Mettl1-KO (*$Mettl1^{KO1}$*) and rescue strain (*$Mettl1^{KO1}$; [Mettl1]*) adult females. Scale bar is 100 µm. Significant differences calculated by two-tailed Student's t-test were indicated on graph.

**Supplementary Figure S2. Loss of Mettl1 didn't affect maintenance of somatic cells of testis**

**(a-c)** Representative immunofluorescence image of *Drosophila* germ or soma cells in WT (*yw*) and Mettl1-KO (*$Mettl1^{KO1}$*) testes. **(a)** Aubergine (Aub) expression in germ cell. **(b)**

Boule expression and localization in 32-cyst (Upper) and 64-cyst (Bottom) cells. **(c)** Upper images indicate hub cell distribution (Fas3) at the apical tip of testes. Bottom images indicate somatic cyst cells (Tj). **(c)** Western blot validation of Mettl1 expression in male carcasses.

**Supplementary Figure S3. Mettl1 methylates target tRNA in a similar manner to that of Mettl1 in other eukaryotes.**

**(a)** Immunoprecipitation from the *attp40[Mettl1-Flag]* ovaries and volcano plots showing enrichment rates and significance levels of each protein as log2 fold change (anti-Flag M2/negative control: non-immune IgG) versus negative log 10 of the student's t-test p. value, and comparison of immunoprecipitates obtained from two different buffer conditions (Left: HEPES-NP40 buffer. Right: Tris-Triton buffer). Blue and red dots represent Mettl1 and Mettl1-interactors, respectively. **(b)** Immunoprecipitation from the *attp40[Mettl1-Flag]* or Oregon R wild-type ovaries (negative control) and volcano plots showing enrichment rates and significance levels of each protein as log2 fold change (*attp40[Mettl1-Flag]*/negative control: Oregon R) versus negative log 10 of the student's t-test p. value, and comparison of immunoprecipitates obtained from two different buffer conditions (Left: HEPES-NP40 buffer. Right: Tris-Triton buffer). Blue and red dots represent Mettl1 and Mettl1-interactors, respectively. **(c)** Left: sequences of tRNA fragments used in $m^7G$ methylation assay. Right: In vitro $m^7G$ methylation assay using recombinant *Drosophila* Mettl1/Wh, tRNA TrpCCA fragment and tRNA TrpCCA fragment (46th G>C). The $m^7G$ containing tRNAs were labelled with $^{14}C$. Toluidine blue *O* staining was done to visualize RNAs.

**Supplementary Figure S4. *Drosophila* Mettl1 catalyzes m$^7$G modification of tRNA in ovaries**

**(a)** Northern blot of the chemical treated total RNAs from control (*Mettl1$^{KO1}$/FM7*) and Mettl1-KO (*Mettl1$^{KO1}$*) ovaries. The probe was designed at around 3′ end of TrpCCA (top) and ValCAC (bottom). **(b)** Schematic diagram of method to calculate cleavage score in TRAC-seq (top). Images of cleavage score of indicated tRNAs (bottom). **(c)** List of RNAs which have Mettl1-dependent m$^7$G modification in ovary.

**Supplementary Figure S5. *Drosophila* Metl1 methylates some tRNAs to stabilize expression but not be involved in regulation *let-7* miRNA expression.**

**(a)** List of three tRNA group defined in this study. (Rules for classifying tRNA was indicated in Fig. 5) **(b)** Northern blot comparison of let-7 miRNA expression between control (*FM7*) and Mettl1-KO (*Mettl1$^{KO1}$*) testes. 2S rRNA is a loading control. **(c)** Comparisons of miRNA let-7 structures between mouse and *D. melanogaster*. Red letters indicate guanosines predicted to be involved in the formation of the quadruplex motif. The asterisk indicates the position of m$^7$G. Gray shading indicates the "RAGGU" motif sequence. The RAGGU motif highlighted in gray is not conserved in *D. melanogaster*.

**Supplementary Figure S6. Identification of ubiquitously expressed and low variance genes (ULVs).**

Schematic diagram of method to obtain ULVs using dual *Drosophila* databases (modENCODE, FlyAtlas2) and compare mRNA expression levels of ULVs between testis and other tissues.

**Supplementary Figure S7. Characterization of ribosome profiling data obtained from fly testes.**

**(a)** Metagene plot for ribosome footprints around start codons. The 5′ end of 29 nt reads were depicted. **(b and c)** Pearson's correlation coefficients (r) ribosome footprints (b) and RNA-Seq reads (c) on each ORFs. The color scales for *r* are represented. (d) Ribosome footprint distribution on the indicated transcript. A-site position of reads are depicted. RPM, reads per million mapped reads.

### Supplementary Figure 1

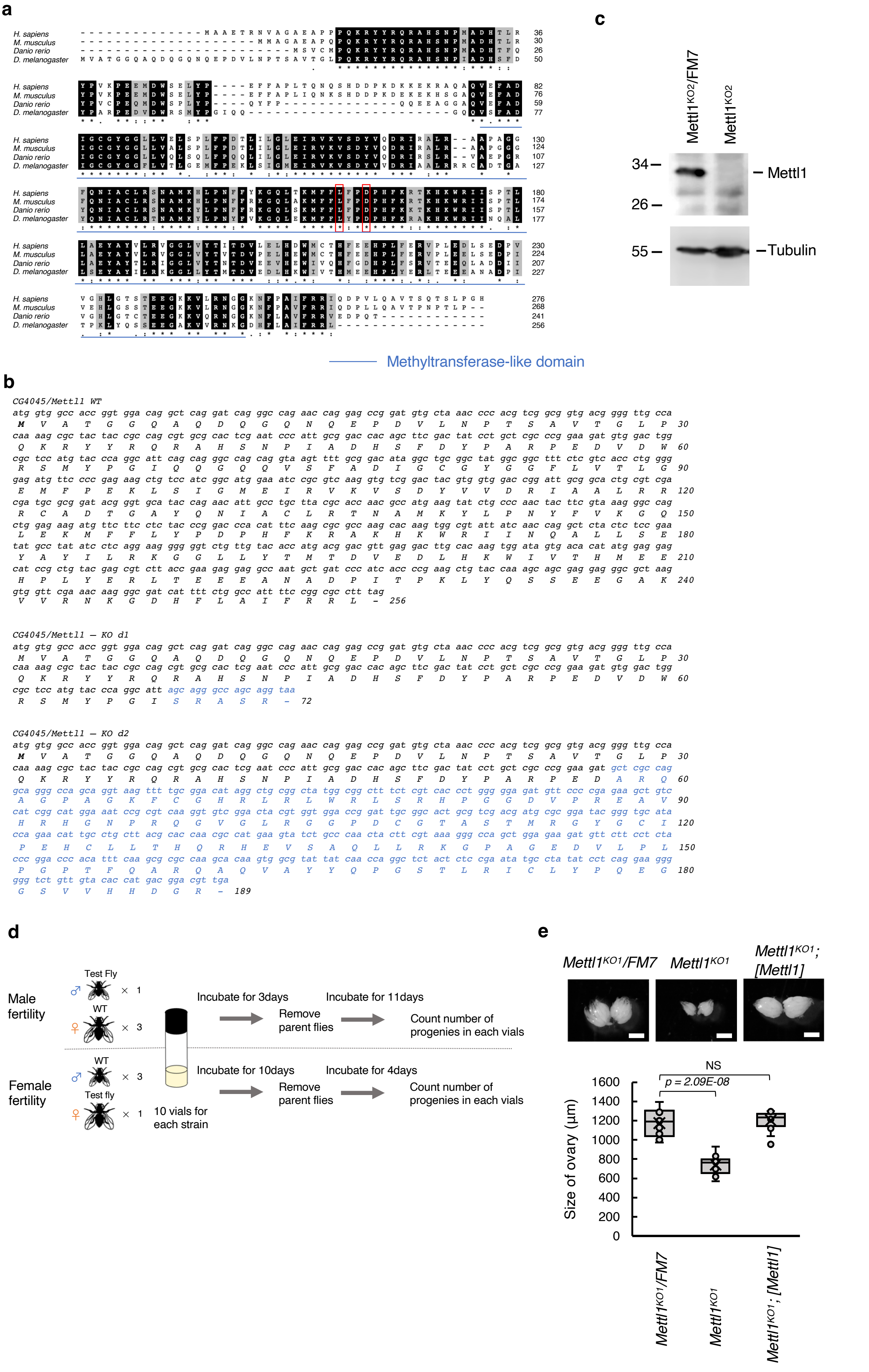

### Supplementary Figure 2

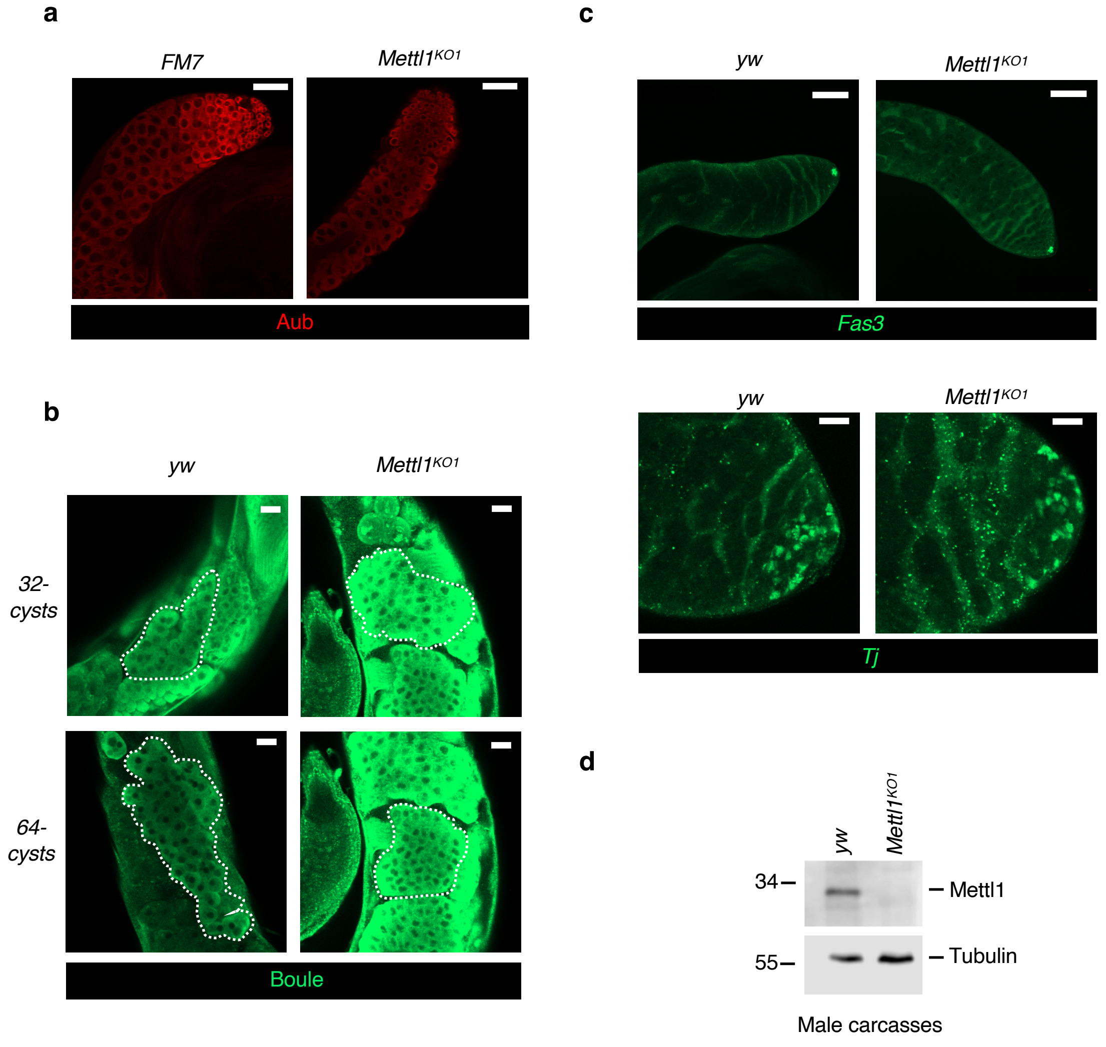

### Supplementary Figure 3

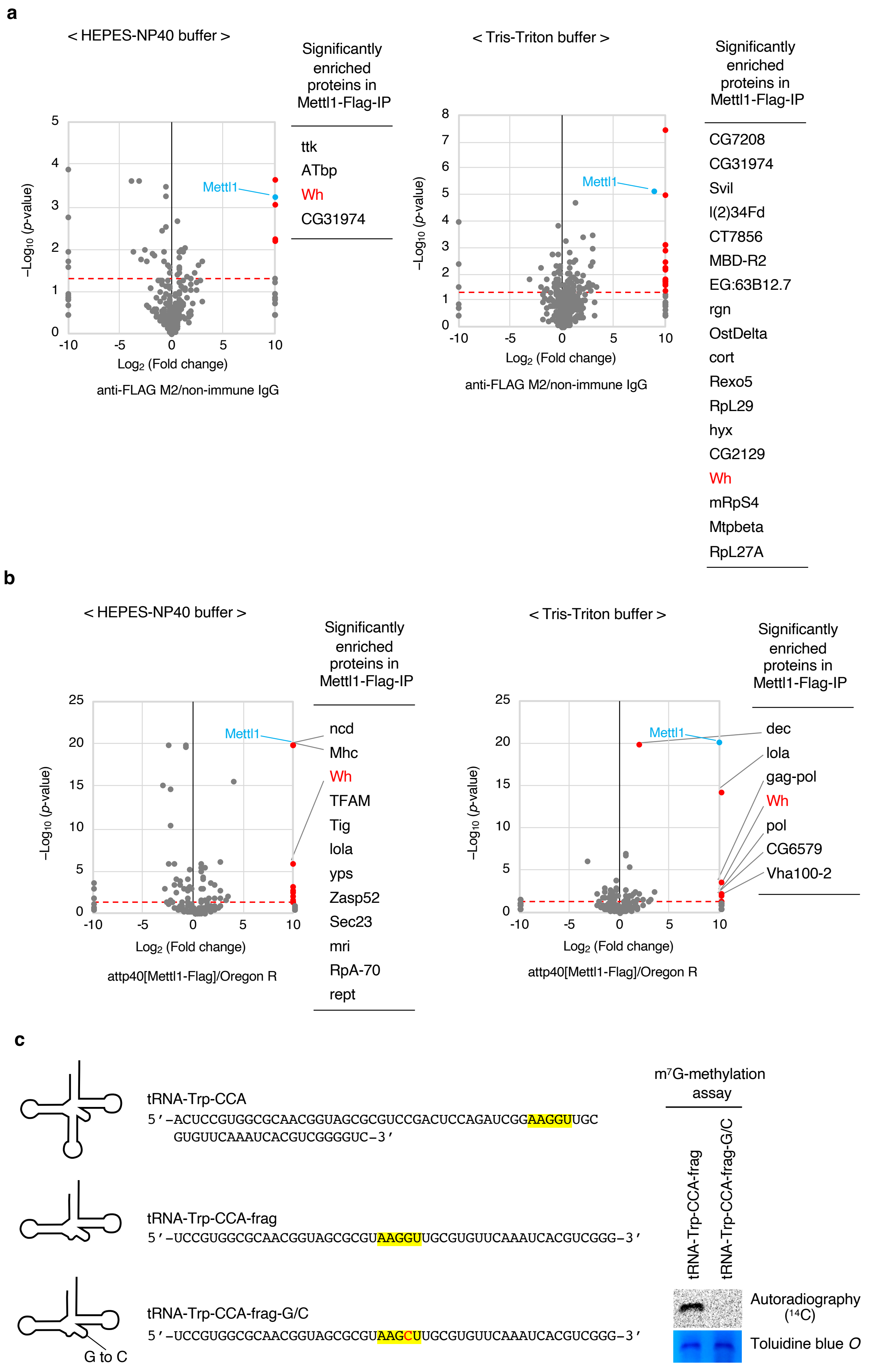

### Supplementary Figure 4

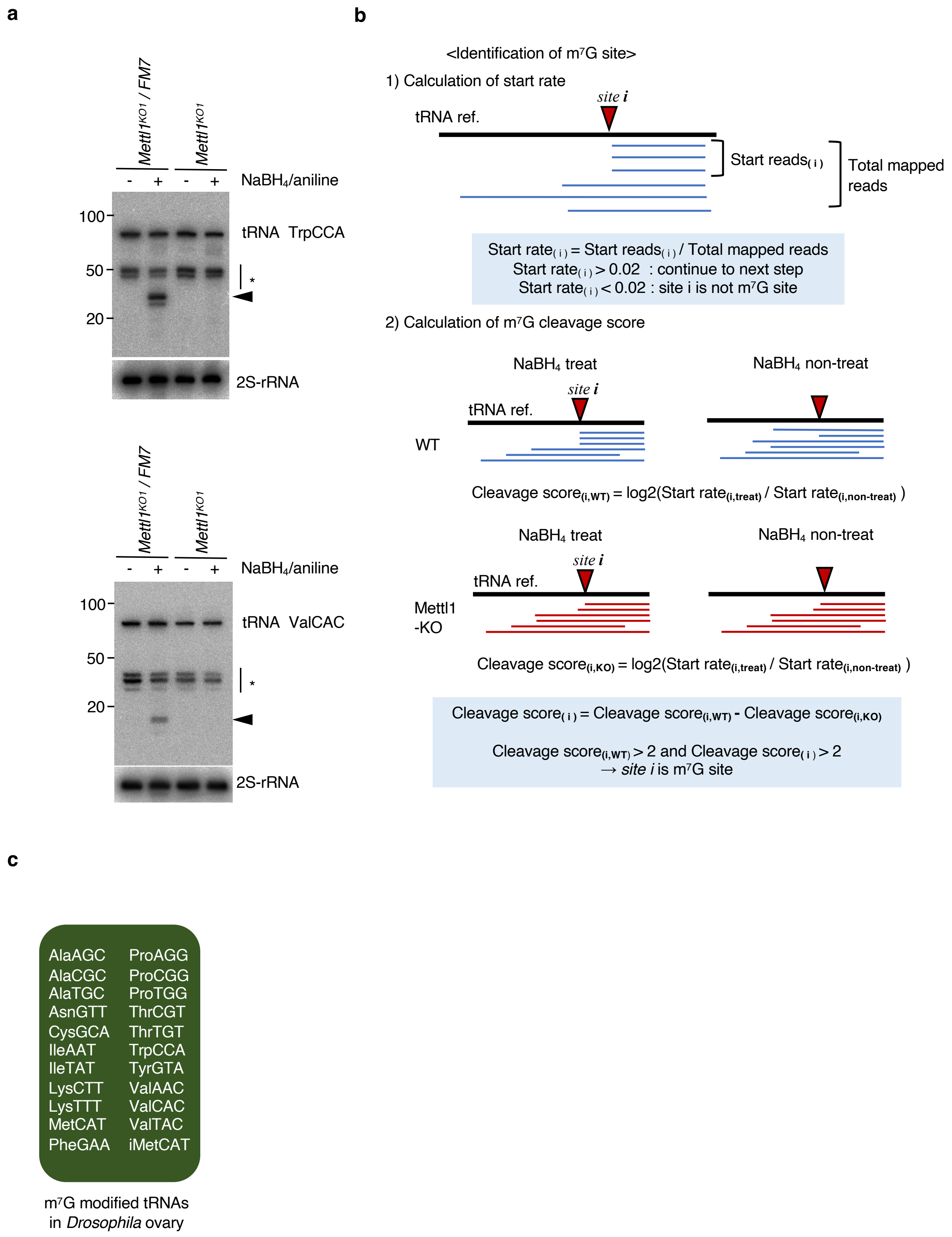

### Supplementary Figure 5

**a**

**Group 1**

AlaAGC, AlaCGC, AlaTGC, ArgTCT, CysGCA, LysCTT, LysTTT, ProAGG, ProCGG, ProTGG, TyrGTA, ValAAC, ValCAC, ValTAC, iMetCAT

**Group 2**

AsnGTT, IleAAT, IleTAT, MetCAT, PheGAA, ThrCGT, ThrTGT, TrpCCA

**Group 3**

ArgACG, ArgCCT, ArgTCG, AspGTC, GlnCTG, GlnTTG, GlnCTC, GluTTC, GlyGCC, GlyTCC, HisGTG, LeuAAG, LeuCAA, LeuCAG, LeuTAA, LeuTAG, SecTCA, SerAGA, SerCGA, SerGCT, SerTGA, ThrAGT

**b**

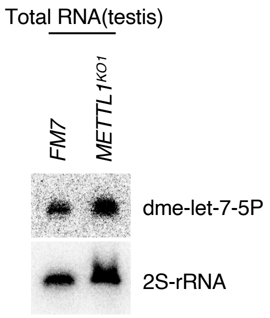


**c**

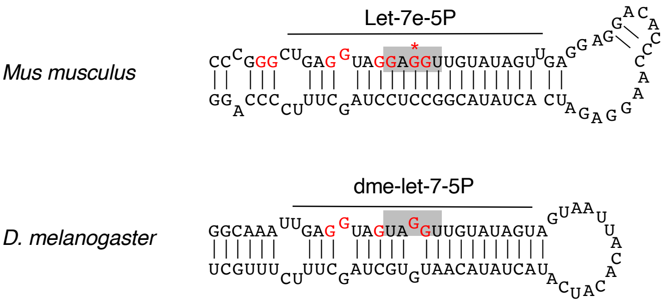

### Supplementary Figure 6

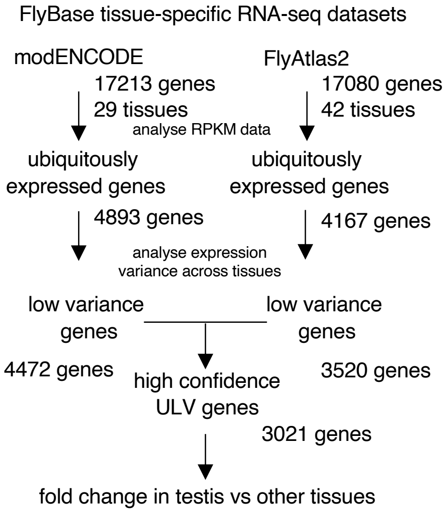

### Supplementary Figure 7

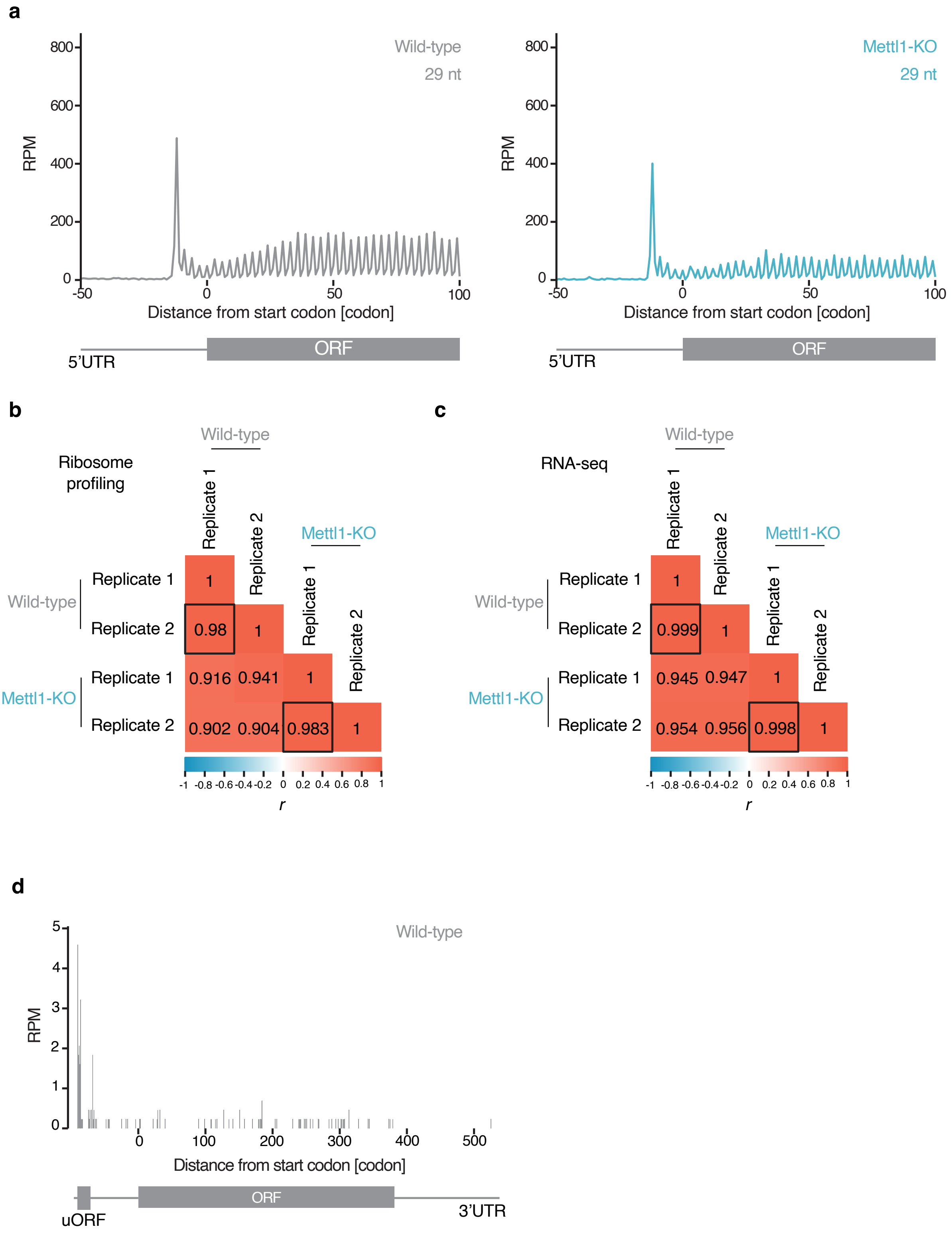

### Supplementary Table 1

List of transcripts analyzed as essential for fertility in the Figure 7d

| Flybase ID | gene |
|---|---|
| FBtr0074950 | ms(3)76 |
| FBtr0083576 | hmw-PA |
| FBtr0081916 | CG8236- |
| FBtr0082559 | CG10014 |
| FBtr0089744 | didum-P |
| FBtr0113086 | loopin- |
| FBtr0087092 | loopin- |
| FBtr0089743 | didum-P |
| FBtr0089745 | didum-P |
| FBtr0306246 | CG6752- |
| FBtr0083029 | CG6752- |
| FBtr0306152 | Lasp-PD |
| FBtr0289982 | Dnah3-P |
| FBtr0087649 | S-Lap7- |
| FBtr0332255 | S-Lap7- |
| FBtr0084382 | bb8-PA |
| FBtr0074936 | Rcd7-PA |
| FBtr0074937 | Rcd7-PB |
| FBtr0113686 | klhl10- |
| FBtr0113685 | klhl10- |
| FBtr0075350 | Lasp-PA |
| FBtr0100145 | Lasp-PB |
| FBtr0306151 | Lasp-PC |
| FBtr0075685 | goddard |
| FBtr0080437 | CG5458- |
| FBtr0300056 | TTLL3B- |
| FBtr0076824 | CG7716- |
| FBtr0301369 | jar-PI |
| FBtr0072161 | CG3121- |
| FBtr0076288 | S-Lap4- |
| FBtr0345266 | jar-PN |
| FBtr0273304 | Duba-PD |
| FBtr0084170 | CG6332- |
| FBtr0084636 | jar-PB |
| FBtr0301367 | jar-PG |
| FBtr0301370 | jar-PJ |
| FBtr0301371 | jar-PK |
| FBtr0334872 | jar-PM |
| FBtr0074948 | ms(3)76 |
| FBtr0301368 | jar-PH |
| FBtr0334871 | jar-PL |
| FBtr0082046 | betaTub-85D |
| FBtr0301787 | Scsbeta ScsβA |
| FBtr0301786 | Scsbeta ScsβA |
| FBtr0081957 | Scsbeta ScsβA |
| FBtr0113206 | Scsbeta ScsβA |
| FBtr0071940 | CG4329- |
| FBtr0334328 | Scsbeta ScsβA |
| FBtr0334327 | Scsbeta ScsβA |
| FBtr0087624 | S-Lap5- |
| FBtr0083636 | CG14305 |
| FBtr0305956 | CG14305 |
| FBtr0076947 | CG32392 |
| FBtr0076948 | CG32392 |
| FBtr0080448 | Prosalp |

### Supplementary Table 2

List of fly strains used in this study

| | Genotype | Experiments |
|---|---|---|
| yw | $y^1$ $w^{1118}$/ Y | WB (Fig.1c, Supp. Fig. 2d), Fertility assay (Fig.1d, f, 2g, 3f, 6b), Immunostaining (Fig. 2b, d, f, Supp. Fig.2b, c), TRAC-seq (Fig. 4), tRNA expression quantigfication (Fig. 5), m7G site cleavage (Fig. 6c), Quantification of eEF1α mRNA level (Fig. 6e), Ribo-seq (Fig. 7, Supp. Fig. 7) |
| Mettl1$^{KO1}$/Mettl1$^{KO1}$ | $y^1$ $w^{1118}$ Mettl1$^{KO1}$/ $y^1$ $w^{1118}$ Mettl1$^{KO1}$ | WB (Fig. 1b), Fertility assay (Fig.1e), TRAC-seq (Supp. Fig.4c)<br>Measurement of ovary size (Mettl1-KO, Supp. Fig. 1e) m7G site cleavage (Supp. Fig.4a), TRAC-seq (Supp. Fig. 4c) |
| Mettl1$^{KO1}$/FM7 | $y^1$ $w^{1118}$ Mettl1$^{KO1}$/ FM7 Kruppel>GFP | WB (Fig. 1b), Fertility assay (Fig.1e), TRAC-seq (Supp. Fig.4c)<br>Measurement of ovary size (WT, Supp. Fig. 1e) m7G site cleavage (Supp. Fig.4a), TRAC-seq (Supp. Fig. 4c) |
| Mettl1$^{KO1}$/Mettl1$^{KO1}$; attp40{Mettl1}/ + | $y^1$ $w^{1118}$ Mettl1$^{KO1}$/ $y^1$ $w^{1118}$ Mettl1$^{KO1}$ ; attp40{y, v, Mettl1} | WB (Fig. 1b), Fertility assay (Fig.1e)<br>Measurement of ovary size (Rescue, Supp. Fig. 1e) |
| Mettl1$^{KO2}$/Mettl1$^{KO2}$ | $y^1$ $w^{1118}$ Mettl1$^{KO2}$/ $y^1$ $w^{1118}$ Mettl1$^{KO2}$ | WB (Supp. Fig. 1c), Fertility assay (Fig.1e) |
| Mettl1$^{KO2}$/FM7, | $y^1$ $w^{1118}$ Mettl1$^{KO2}$/ FM7 Kruppel>GFP | WB (Supp. Fig. 1c), Fertility assay (Fig.1e) |
| Mettl1$^{KO2}$/Mettl1$^{KO2}$; attp40{Mettl1} | $y^1$ $w^{1118}$ Mettl1$^{KO2}$/ $y^1$ $w^{1118}$ Mettl1$^{KO2}$ ; attp40{y v Mettl1} | Fertility assay (Fig.1e) |
| Mettl1$^{KO1}$, Mettl1$^{KO1}$/Y | $y^1$ $w^{1118}$ Mettl1$^{KO1}$/Y | WB (Fig. 1c, Supp. Fig. 2d) Fertility assay (Fig. 1d, f, 2g, 3f, 6b), Observation of seminal vesicle (Fig. 2c), Immunostaining (Fig. 2d, f, Supp. Fig. 2a, b, c), Observation of bundle structure in testis (Fig. 2e), m7G site cleavage (Fig. 4b, 6c), TRAC-seq (Fig. 4), Quantification of tRNA expression (Fig. 5), Validation of let-7 expression (Supp. Fig. 5b), Quantification of eEF1α mRNA level (Fig. 6e), Ribo-seq (Fig. 7, Supp. Fig. 7) |
| FM7, FM7/ Y | FM7/Y | Observation of seminal vesicle (Fig. 2c), Observation of bundle structure<br>in testis (Fig. 2e), m7G site cleavage (Fig. 4b, c)<br>Validation of tRNA expression (Fig. 5d), Validation of let-7 expression (Supp. Fig. 5b), Immunostaining (Supp. Fig. 2a) |
| Mettl1$^{KO1}$/ Y; attp40{Mettl1}/ + | $y^1$ $w^{1118}$ Mettl1$^{KO1}$/ Y ; attp40{y v Mettl1} | Fertility assay (Fig.1d, 3f), Immunostaining (Fig. 2f) |
| Mettl1$^{KO2}$/ Y | $y^1$ $w^{1118}$ Mettl1$^{KO2}$/ Y | Fertility assay (Fig. 1d) |
| Mettl1$^{KO2}$/ Y; attp40{Mettl1}/ + | $y^1$ $w^{1118}$ Mettl1$^{KO1}$/ Y ; attp40{y v Mettl1} | Fertility assay (Fig. 1d) |
| Mettl1$^{KO1}$/ Y; attp40{nosP-Mettl1}/ + | $y^1$ $w^{1118}$ Mettl1$^{KO1}$/ Y ; attp40{y v nosP-Mettl1} | Fertility assay (Fig.2g) |
| Mettl1$^{KO1}$/ Y; attp40{Mettl1-Cd}/ + | $y^1$ $w^{1118}$ Mettl1$^{KO1}$/ Y ; attp40{y v Mettl1-Cd} | Fertility assay (Fig. 3f) |
| $y^2$ $cho^2$ $v^1$ / Y ; attp40(Mettl1-3 ×FLAG) | $y^2$ $cho^2$ $v^1$ / Y ; attp40(Mettl1-3 ×FLAG) | Immuno staining (Fig. 2b), LC-MS/MS analysis (Fig. 3a, Supp. Fig. 3a) |
| Wh$^{56}$/ Y | w* wuho$^{56}$/ Y | m7G site cleavage (Fig. 4c) |
| FM7c/ Y | FM7c/ Y | m7G site cleavage (Fig. 4c) |
| Mettl1$^{KO1}$/ Y; attp40{UASt-eEF1α}/Act-GAL4} | Mettl1$^{KO1}$/ Y; attp40{y v UASt-eEF1α}/ P{w Act5c-GAL4} | Fertility assay (Fig. 6b) |
| w* / Y ; dj-GFP / CyO | w* / Y ; P{dj-GFP.S}AS1 / CyO | Observation of Dj (Don juan) expression in testis (Fig. 7h) |
| Mettl1$^{KO1}$/ Y; dj-GFP | $y^1$ $w^{1118}$ Mettl1$^{KO1}$/ Y ; P{dj-GFP.S}AS1/ CyO | Observation of Dj (Don juan) expression in testis (Fig. 7h) |

### Supplementary Table 3

The sequence of gRNA for Mettl1-KO and nucleotides for sanger sequence

| No. | Sequence (5' - 3') | Use |
|---|---|---|
| gRNA Mettl1 | ATGTACCCAGGCATTCAGCA | gRNA for Mettl1-KO |
| Mettl1 Fw | CAGCCCTGCTGCTAACTAGG GTCCACCACGTAGTCCGACACCTTGACG | Sanger sequence for checking Mettl1 deletion |
| Mettl1 Rv | ATATCGTTTGAATGCCCCCAACGCAAGC | Sanger sequence for checking Mettl1 deletion |

The probe sequence for northern blot analysis

| Gene | Sequence (5' - 3') | Use |
|---|---|---|
| 2S-rRNA | TACAACCCTCAACCATATGTAGTCCAAGCA | detection of 2S-rRNA |
| mir-let-7-5P | ACTATACAACCTACTACCTCA | detection of mir-let-7-5P |
| tRNA LeuAAG/TAG | CAGCGGTGGGATTCGAACC | Detection of tRNA LeuAAG/TAG |
| tRNA ProTGG | CTCAACCGGGATTTGAACCC | Detection of tRNA ProTGG |
| tRNA TrpCCA | TGACCCCGACGTGATTTG | Detection of tRNA TrpCCA |
| tRNA ValCAC | GTTTTCGCCCGGGTTCGAACC | Detection of tRNA ValCAC |

The sequence of synthesized tRNA fragment used in *in vitro* methylation assay

| Gene | Sequence (5' - 3') | Use |
|---|---|---|
| tRNA (TrpCCA) | ACUCCGUGGCGCAACGGUAGCGCGUCCGACUCCA GAUCGGAAGGUUGCGUGUUCAAAUCACGUCGGGG UC | *in vitro* methylation assay |
| tRNA TrpCCA-frag | UCCGUGGCGCAACGGUAGCGCGUAAGGUUGCGU GUUCAAAUCACGUCGGG | *in vitro* methylation assay |
| tRNA TrpCCA-frag-G/C | UCCGUGGCGCAACGGUAGCGCGUAAGCUUGCGU GUUCAAAUCACGUCGGG | *in vitro* methylation assay |

The sequence of primer for vector construction

| Gene | Sequence (5' - 3') | Use |
|---|---|---|
| Mettl1 Fw | ATCGAATTCCCTGCAGGCCTGGAATATCAGGAACG TGAGCACCG | Amplification of Mettl1 genomic region |
| Mettl1 Rv | GCGTGACGCGCGGCCGCCTGACCAATCACACGCG CACCGCCATC | Amplification of Mettl1 genomic region |
| Mettl1-Cd Fw | GATGTTCTTCgcCTACCCGGcCCCACATTTCAAGCG CGCCAAGC | Inducing mutation to Mettl1-transgene |
| Mettl1-Cd Rv | TGAAATGTGGGgCCGGGTAGgcGAAGAACATCTTCT CCAGCTGGCCCT | Inducing mutation to Mettl1-transgene |
| eEF1α1 Fw | GGGGCGCGCCGCTAGCATGGGCAAGGAAAAGATT CACAT | Amplification of eEF1α1 CDS |
| eEF1α1 Rv | GCGTGACGCGCGGCCGCCTACTTCTTGCCCTTGGT GG | Amplification of eEF1α1 CDS |
| Mettl1 (CDS) Fw | acactacgtagaattcGTGGCCACCGGTGGACAGGC | Amplification of Mettl1 CDS |
| Mettl1 (CDS) Rv | tagaggatccgaattcCTAAAGGCGCCGGAAAATGGCCAG | Amplification of Mettl1 CDS |
| Wh (CDS) | tgggatccccGAATTCTGCACAACAATTTCGTTCGCGGA AC | Amplification of Wh CDS |
| Wh (CDS) | gatgcggccgCTCGAGTTAGCCGCACTTCTGCTGCTGC | Amplification of Wh CDS |
